# Structures of wild-type and selected CMT1X mutant connexin 32 gap junction channels and hemichannels

**DOI:** 10.1101/2023.03.08.531661

**Authors:** Chao Qi, Pia Lavriha, Erva Bayraktar, Anand Vaithia, Dina Schuster, Micaela Pannella, Valentina Sala, Paola Picotti, Mario Bortolozzi, Volodymyr M. Korkhov

## Abstract

In myelinating Schwann cells, communication between myelin layers is mediated by gap junction channels (GJC) formed by docked connexin 32 hemichannels (HCs). Mutations in Cx32 cause the X-linked Charcot–Marie–Tooth disease (CMT1X), a degenerative neuropathy with no cure. A molecular link between Cx32 dysfunction and CMT1X pathogenesis is still missing. Here, we describe the high resolution cryo-EM structures of the Cx32 GJC and HC, along with two CMT1X-linked mutants, W3S and R22G. While the structures of wild-type and mutant GJCs are virtually identical, the HCs show a major difference: in the W3S and R22G mutant HCs, the N-terminal helix partially occludes the pore, consistent with an impaired HC activity. Our results suggest that HC dysfunction may be involved in the pathogenesis of CMT1X.

**One-Sentence Summary:** Connexin 32 channel structures reveal a gating helix defect in CMT1X disease-associated mutant hemichannels

## Introduction

Connexin-mediated communication is one of the major pathways of intercellular signaling (*1, 2*), involving complex physiological and pathological processes, such as electrical activity of the heart (*3*), neuronal signaling (*4*), release of hormones (*5*), immunity (*6*), inflammation (*7*), cancer (*8*), and cell death (*9*). The importance of connexin-mediated communication is underscored by at least 28 genetic diseases linked to mutations in the 21 genes encoding connexins in humans (*10*), (*11*).

Connexin isoforms share a conserved molecular architecture: each is a 4-transmembrane (TM) domain protein that assembles into hexamers called connexons or hemichannels (HCs). Two HCs expressed at juxtaposed plasma membrane regions of adjacent cells, or even the same cell, can dock together to form a GJ channel (GJC) (*11*). Tens to thousands of GJCs assemble in a regular hexagonal pattern (a GJ plaque) which allow direct intercellular exchange of ions, metabolites, second messengers (e.g., cAMP, IP3) or peptides (*12*). Unlike GJCs, connexin HCs are normally closed at rest but can open under physiological conditions, allowing sustained ion fluxes and release of ATP, NAD^+^, glutamate and other signalling molecules into the extracellular space (*13*).

Connexin-32 (Cx32), encoded by the *GJB1* gene, is strongly expressed in the liver (*14*), but also found in various tissues throughout the body, including the central and peripheral nervous systems (*15, 16*). In the peripheral nervous system, Cx32 is expressed in myelinating Schwann cells, particularly in non-compact regions of the myelin sheath (*15, 17*), where GJCs were proposed to provide a radial diffusion pathway between the abaxonal and adaxonal regions (**Fig. 1A**) (*18*). Moreover, extracellular release of ATP by Cx32 HCs has been proposed to support purinergic signalling triggered by neuronal activity, thus regulating Schwann cell myelin maintenance (*19, 20*).

**Fig. 1.**
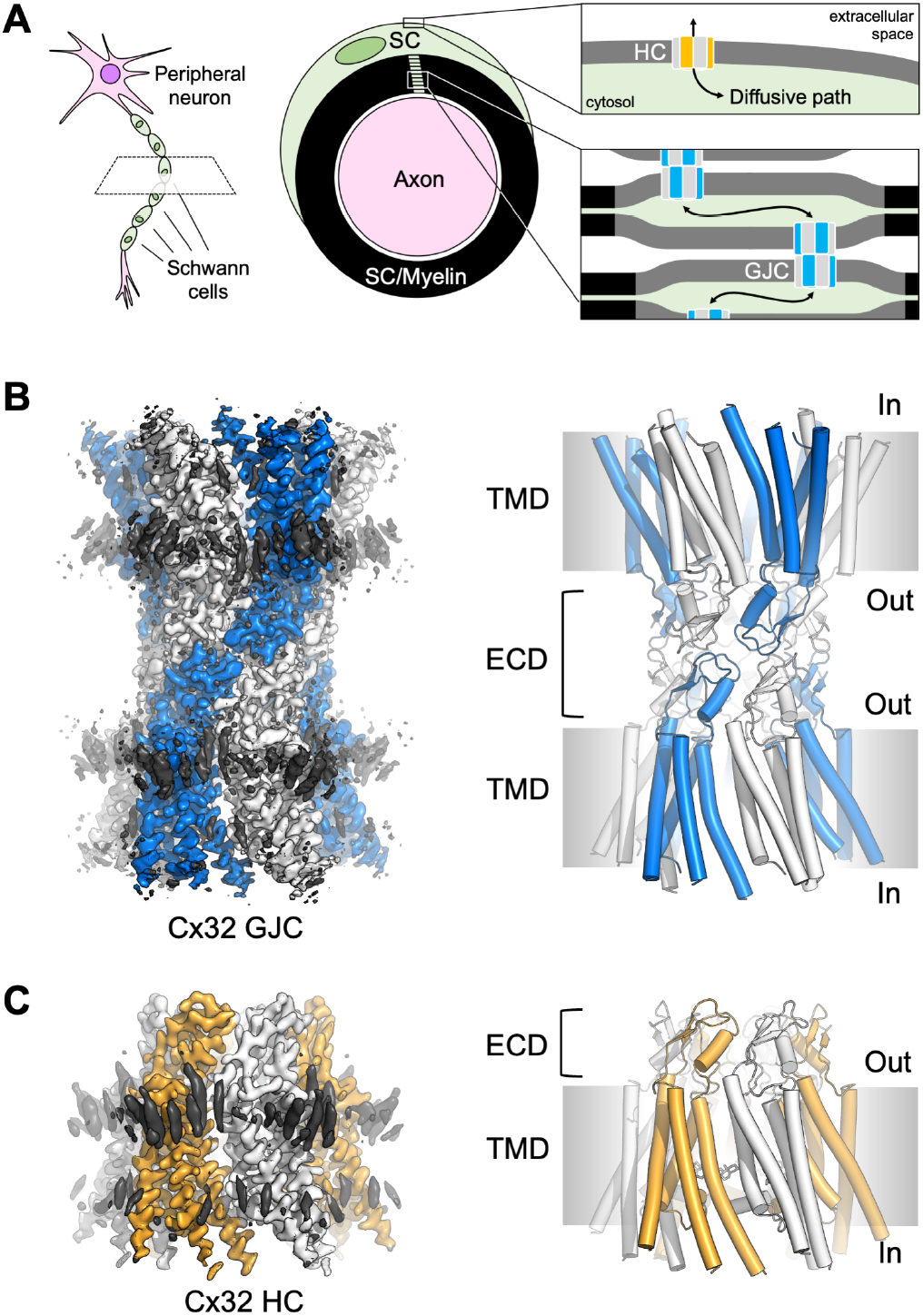
Cryo-EM structures of Cx32 GJC and HC. (**A**), An illustration of Cx32 GJCs and HCs in distinct membrane compartments of the myelinating Schwann cells (“SC”). (**B**) Cryo-EM map and model of Cx32 GJC. Cx32 is surrounded by lipid like molecules at outer leaflet and inner leaflet of the membrane (black density). (**C**) Cryo-EM map and model of Cx32 HC.

Mutations of Cx32 that affect its biosynthesis, folding, assembly, trafficking or channel properties are the leading cause of the X-linked form of Charcot-Marie-Tooth disease (CMT1X), a neuro-muscular degenerative disorder (*21-23*). Axonal and myelin defects in peripheral neurons of CMT1X patients lead to progressive sensory abnormalities and muscle atrophy. The disease occurs at any stage of a patient’s life and has currently no cure (*20, 24*). Loss of function of GJCs and/or HCs due to Cx32 mutations is presumed to be the prevailing cause of the disease (reviewed in detail in (*20*)), although the molecular and cellular mechanisms underlying its pathogenesis remain unclear.

Despite the importance of Cx32 in human physiology and pathology, the structure of this protein has not yet been described. Insights into the structure-function relationships of Cx32 have been derived indirectly through homology modeling from the structures of other connexin proteins (*25-27*).

To date, the structures of four connexin complexes have been experimentally determined: Cx26 GJC (*28-31*), Cx46/50 GJC (*32, 33*), Cx31.3 HC (*34*) and Cx43 GJCs and HC (*35*). These homologous structures have provided important clues to the shared architectural features and assembly of connexin-mediated GJCs and HCs, but also revealed substantial differences between them. To advance our understanding of the molecular basis of Cx32 channel activity and regulation in health and disease, we utilized cryo-electron microscopy (cryo-EM) to determine the structures of the HC and GJC formed by Cx32 WT as well as the two mutants, W3S and R22G, linked to CMT1X.

## Results

### Structures of the Cx32 GJC and HC

The Cx32 protein tagged with a C-terminal YFP was chosen for large-scale protein expression for structural studies. To ascertain that this construct can be used to generate active GJCs, we used the dual-patch-clamp technique to measure the GJ conductance (*g*_j_) in HeLa DH cells (Fig. S1). The measured *g*_j_ values confirmed the expression of functional Cx32 GJCs, whereas control cells exhibited negligible intercellular communication. Both Cx32 GJCs (*g*_j_ values) and HCs (*g*_m_ values) were sensitive to cytoplasm acidification mediated by extracellular CO_2_, consistent with our previous studies (*27*).

Cx32-YFP construct was expressed in HEK293F cells, and the protein preparations were purified using anti-GFP nanobody affinity chromatography, followed by size-exclusion chromatography (Fig. S2A). The protein samples with the affinity tag removed were concentrated, vitrified on cryo-EM grids and used for cryo-EM analysis (Fig. S3, S6A). The sample featured very well preserved full GJC structures, evident particularly in the thin ice areas of the cryo-EM grids (Fig. S3). We determined the cryo-EM reconstruction of the Cx32 GJC in D6 symmetry, at 2.14 Å resolution, revealing the fine details of the protein structure (**Fig.1B-C**, Figs. S3C, S7A). The quality of the resulting density map revealed alternative side-chain conformations, ordered water molecules (Fig. S7A, C), and a number of lipid- or detergent-like density elements at the protein-lipid domain boundary (**Fig. 1B**). Furthermore, using the same micrographs and focusing on processing of the smaller HC particles, we were able to reconstruct the structure of the Cx32 HC at 3.06 Å resolution in C6 symmetry (**Fig. 1C**, Figs. S3C, S6B). Processing of the cryo-EM data in C1 symmetry confirmed the presence of all density map features observed with C6 symmetry imposed (Fig. S3C; Table S1).

### Comparison of Cx32 GJC and HC structures

The overall conformation of the Cx32 HC appears to be very similar to the connexon of a fully assembled GJC (**Fig. 2A-D**, Fig. S8A). There are two key differences that concern the extracellular loops (ECL1 and ECL2) and the N-terminal helix (NTH) regions of Cx32.

**Fig. 2.**
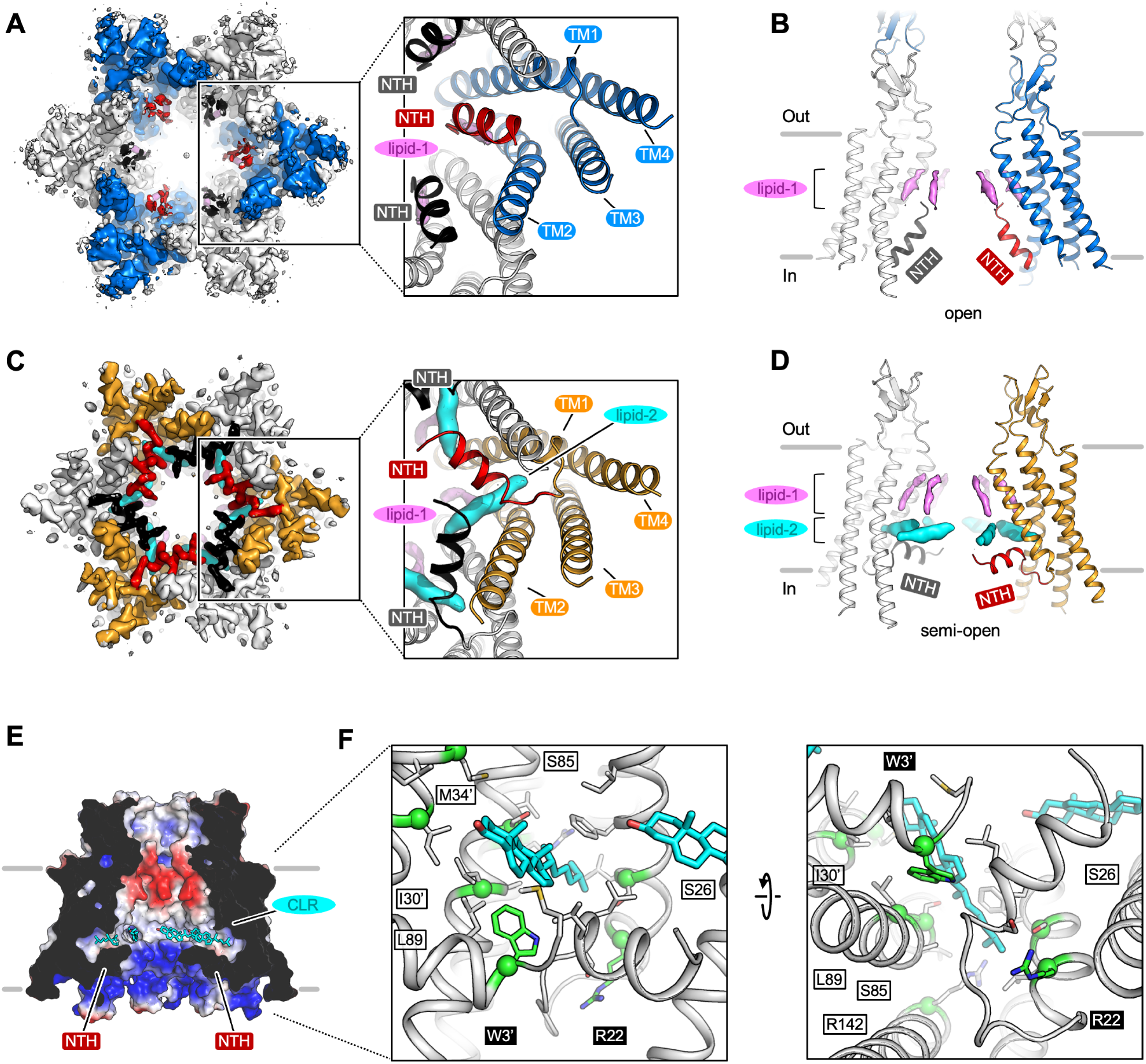
Comparisons of Cx32 GJC and HC. (**A-B**) Cryo-EM map (bottom view) and model (side view) of Cx32 GJC. The NTH (red and black) inserts to the channel pore, representing the open state of GJC. The lipid1 density is labeled using pink color. (**C-D**) Cx32 HC map (bottom view) and model (side view). The NTH (red and black) is parallel to the membrane layer, shrinking the pore of Cx32 HC to about 11Å. Lipid2 (cyan) is stabilized by NTH and TM1, TM2. (**E**) Electrostatic map of Cx32 HC. Cholesterol (CLR) molecules (cyan) are fitted to the lipid2 position. (**F**) Detailed view of the interaction between CLR and NTH, TM1, TM2. All the residues within 4Å to CLR are shown as sticks. Two important residues, W3 and R22, are shown as green color.

The ECL1 and ECL2 of the Cx32 GJC connexon show a slight conformational rearrangement, compared to those in the HC (Fig. S8A, B). This conformational rearrangement is very similar to that of Cx43 we observed previously (*35*). Cx32 belongs to the β group connexins based on the inter-HC interface. It shares the same interface residues with Cx26 (N54-N57 in ECL1 and N175-D178 in ECL2, Fig. S8C). As a consequence of this similarity in sequence and structure, Cx32 is capable of forming heterotypic GJCs with Cx26. In contrast, the α group of connexins, which includes Cx43, is not compatible with the β group. Nevertheless, Cx32 and Cx43 utilize a similar mechanism to assemble a GJC, whereby ECL1 and ECL2 move outward to dock the two HCs and form a full GJC.

The NTH of the HC is very well ordered and adopts a previously undescribed conformation (**Fig. 2C-D**), unlike the NTH of the GJC that appears to be poorly ordered. We did not build the NTH in our Cx32 GJC model due to the poor quality of the GJC density map in the corresponding region; only approximate placement of the NTH is possible (**Fig. 2A-B**). Based on the mass spectrometric analysis, the Met1 residue is present in our Cx32 sample, which is different from Cx31.3 (*34*) and Cx43 (*35*) (Fig. S9), and therefore our model of the Cx32 HC includes Met1. The NTH of Cx32 HC is positioned parallel to the membrane plane, maintaining an open pore conformation (**Fig. 2C-D**). Although the structures of both the GJC and the HC represent an open pore state, the rigid conformation of the NTH in the Cx32 HC constricts the pore to a diameter of ∼11 Å at the cytosolic side (**Fig. 2B**). In contrast, the pore diameter in the GJC, based on the ordered region of the protein, is ∼15 Å (Fig. S10C), not accounting for the present flexible NTH regions. The lack of clearly defined NTH conformations in the GJC suggests that the two connexons are fully open and may allow free unselective movement of both anions and cations (*36*), as well as small molecules. The arrangement of the NTH in the HC not only reduces the size of the pore, but also dramatically remodels the charge distribution at the cytosolic face of the channel. The strong positive charge at the cytosolic entrance into the Cx32 HC pore may play a role in ion selectivity or gating of the channel, although it is important to note that the channel remains open in this conformation (**Fig. 2E**, Fig. S10C).

### Comparison of the Cx32 GJC and HC to other connexin structures

The overall conformation of the Cx32 GJC is similar to the conformations observed in the structures of GJCs formed by Cx26, Cx46/50 and Cx43 (Fig. S11A). The main distinction between these channel structures is the NTH region. The NTH domains of Cx26 and Cx46/50 are structured and point deeply into the pore. The NTH of Cx43 points towards the center of the pore, closing the gate. The NTH of Cx32 is flexible and we only observe a weak density at the position corresponding to the NTH of Cx26 and Cx46/50 (**Fig. 2A**, Fig. S11A).

The recently determined structure of the Cx31.3 HC revealed a “semi-closed” conformation of the gate, wherein the NTH points to the middle of the pore (the diameter is about 8 Å) (Fig. S11B), similar to the Cx43 GJC. Compared to Cx31.3 and Cx43, the NTH of Cx32 HC arranges itself parallel to the membrane plane but pointing towards the adjacent connexin subunit. This conformation is reminiscent of an iris-like structure in the hexameric NTH arrangement (**Fig. 2C-D**).

### Lipid decoration of Cx32 GJC and HC

As has been observed in the structures of Cx31.3, Cx46/50 and Cx43 (*33-35*), the Cx32 maps (GJC and HC) feature lipid-like densities at the protein-lipid interface (**Fig. 1B-C**). Our Cx32 protein was solubilized from the HEK293 cell membranes using a mixture of N-dodecyl-β-D-maltopyranoside (DDM) and a cholesterol analogue cholesteryl hemisuccinate (CHS), with subsequent detergent exchange to digitonin, as detailed in Material and Methods. Based on the shapes of the observed densities, we interpreted them as protein-bound CHS molecules. However, it is likely that these binding sites in proximity of the hydrophobic surfaces of the protein may be occupied either by detergent molecules, or by cholesterol, phospholipids or other lipid-like molecules from the native membrane environment of Cx32. This possibility is supported by the recent structure of GJC formed by Cx46/50, which was determined in the presence of phospholipids bound at specific sites to the hydrophobic surface of Cx46/50 (*33*). The same is also true for our Cx43 GJC structure in lipidic environment (*35*). The lipids surrounding the GJCs and HCs may be important for channel assembly, stability and function.

Interestingly, we also observed two lipid densities in the interior of the Cx32 HC, which we refer to as “lipid-1” and “lipid-2” (**Fig. 2B, D**). Lipid-1 is aligned along the pore close to the TM1 and TM2 of both the Cx32 GJC and HC (**Fig. 2B, D**), perpendicular to the membrane plane. A similar lipid density has been observed in the HC structure of Cx31.3, where it was interpreted as a LMNG molecule, suggesting it as a conserved lipid binding site. Likewise, our Cx43 structures in detergent and in nanodisc also revealed the presence of a very similar lipid molecule in the pore (*35*). Lipid-2 inserts to a pocket (NT pocket) formed by TM1, TM2 and NTH (**Fig. 2D-F**). We interpret lipid-2 as a bound cholesterol molecule based on a fit of cholesterol to the density (Fig. S7B); CHS and digitonin match the density less well. The NT pocket and the bound lipid-2 molecule (**Fig. 2D-F**) are specific to the Cx32 HC. The NTH is not ordered in the GJC reconstruction and the density element corresponding to lipid-2 is missing (**Fig. 2A-D**).

### Structure-based insights into the pathogenesis of CMT1X

Multiple mutations in Cx32 have been found to be associated with CMT1X, and we can now locate these mutations in the experimentally determined Cx32 structures (Fig. S12). Mapping of the disease-linked mutations with known molecular phenotypes can help us in understanding the mechanism behind their deleterious effects in patients suffering from CMT1X. The Cx32 mutations can be grouped on the basis of defects related to: (i) assembly and cell surface trafficking of Cx32 HCs (Fig. S11, cyan), (ii) formation of GJCs (yellow), (iii) gating mechanisms of the assembled HCs and GJCs (orange) (*20*). Although the defects in folding, assembly, and trafficking in certain mutants of Cx32 are difficult to explain based on the structures, our 3D reconstruction allow us to explain the effects of the mutations proximal to the NTH or the NT pocket. We selected two CMT1X mutations proximal to the NTH and the NT pocket, W3S and R22G, to characterize their effect structurally and functionally. Our structures show that these two residues engage in many interactions that stabilize the NT pocket and thus likely participate in the molecular gating. The two mutants of Cx32, W3S (*37*) and R22G (*38*) have been reported to be capable of assembling the connexon and reaching the plasma membrane, without participating in functional GJs when expressed in HeLa cells and *Xenopus* oocytes, respectively. Similar to the WT Cx32, we purified the mutated proteins (Fig. S3B, C) and determined their structures using single particle cryo-EM (Figs. S4-5, S6C-F). The GJC structures of W3S and R22G were determined at 2.63 Å and 2.48 Å resolution, respectively. In addition to the 3D reconstructions of the mutant GJCs, we determined the structures of W3S and R22G HCs at 2.99 Å and 3.67 Å resolution (Table S1).

Comparison of the two mutant GJC structures to the WT GJC did not reveal any obvious differences (Fig. S10A): both mutant channels showed near-identical features of the NTH density, consistent with the open state. In contrast, the W3S and R22G HCs showed stark differences to the WT Cx32 HC in the NTH region (**Fig. 3A-C**, Fig. S10B). The lipid-2 is no longer present in the HC density maps of W3S and R22G and, unlike the GJC, the NTH of each of the mutants is displaced closer to the center of the pore, resulting in constriction of the central gate. The diameter of the central pore for both mutants is ∼6 Å, compared to ∼11 Å in the case of the WT HC (**Fig. 3A-B**). Thus, in both the selected W3S and R22G mutants, the HCs show a substantial closure of the NTH gate with respect to the WT channel, while the mutant GJCs remain open based on their 3D reconstructions.

**Fig. 3.**
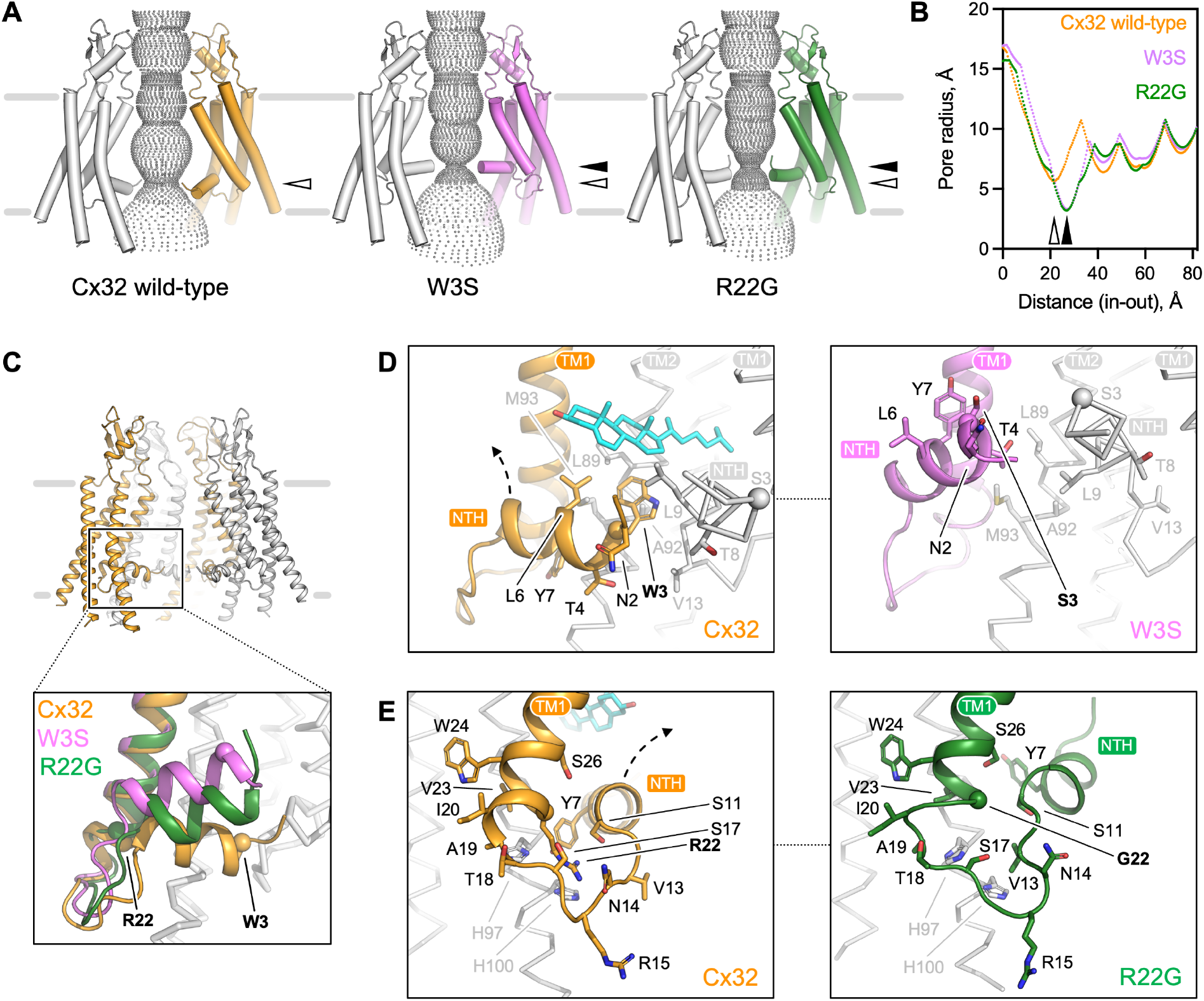
Comparison of the WT Cx32 HC structure to the cryo-EM structures of W3S and R22G Cx32 mutants. (**A**) Substrate conduction pathways for Cx32, W3S and R22G HCs, calculated using Hole software. The arrow indicates the narrowest part of the path. Hollow arrow for Cx32 and black arrow for mutants. (**B**) The pore radius along the substate conduction pathway was calculated using hole in **A**. (**C**) Structural alignment of NTH between Cx32 HC (orange), W3S HC (pink) and R22G HC (green). Key residues, W3 and R22, are labeled as spheres. (**D-E**) The detailed view of NTH in Cx32, W3S and R22G. The key residues within 4 Å close to CLR are shown as sticks. The pink and green arrow indicate the movement of NTH in W3S and R22G compared to Cx32 HC.

### Surface expression and HC dysfunction of the W3S and R22G mutants

To validate the structural observations, we performed functional assays in HEK293F and HeLa DH cells to study the permeability and gating of WT and mutant Cx32 channels. For this purpose, we used a construct containing connexin sequence followed by an internal ribosome entry site (IRES) and a cytosolic yellow fluorescent protein (YFP) to identify the transfected cells. Immunocytochemistry (ICC) experiments in both cell lines revealed that WT Cx32, W3S and R22G localized to the plasma membrane and formed GJ plaques, observed as fluorescent lines on the plasma membrane between two cells (Fig. S13-14). Consistent with this observation, quantification of relative plasma membrane protein expression using surface protein biotinylation followed by mass spectrometric analysis showed that the levels of cell surface expression of the WT Cx32 and the two mutants were comparable (Fig. S15). Interestingly, despite the ability of the mutants to reach the cell surface in both HEK293F and in HeLa DH cells, the properties of the GJ plaques formed by W3S and R22G in these two cell types were dramatically different. In HEK293F cells the GJ plaques formed by the mutants were indistinguishable from those formed by the WT Cx32 (Fig. S13E-F), whereas in HeLa DH cells both mutants produced fewer and smaller GJ plaques (Fig. S13K-L).

To assess the permeability properties of WT and mutant GJ plaques, we performed patch-clamp and Fluorescence Recovery After Photobleaching (FRAP) experiments in both HEK293F and HeLa DH cells. Although HEK293 cells have previously been reported to endogenously express Cx43 and Cx45 (*39, 40*), with surface biotinylation followed by mass spectrometric (MS) analysis, we could confirm that under our conditions of overexpression Cx32 is the dominant connexin species in the plasma membrane of HEK293F cells. This allowed us to use these cells as a model system. In HEK293F cells, dual patch-clamp experiments indicated that the cells expressing WT Cx32 are significantly stronger coupled than control cells (**Fig. 4A** and Fig. S16). The mutant GJs also showed a trend for higher electrical coupling compared to the control cells. Indeed, we expect that the *g*_*j*_ values derived for both WT and mutant GJCs are higher in reality, since only *g*_j_ values ranging lower than 10-15 nS in our measurements minimize the need for corrective methods based on electrode series and cellular input resistances (*41*). The same dual patch-clamp protocol performed in HeLa DH cells revealed a clear difference between the WT Cx32 and the mutants: essentially no coupling could be observed for the latter (Fig. S16A-C). This is likely the result of the deficient GJ plaque formation ability of the mutants in HeLa DH cells, as revealed by our ICC experiments (Fig. S13). This finding highlights the dependence of GJ activity assays on the cell type and the importance of validating GJ activity data using more than one cell system.

**Fig. 4.**
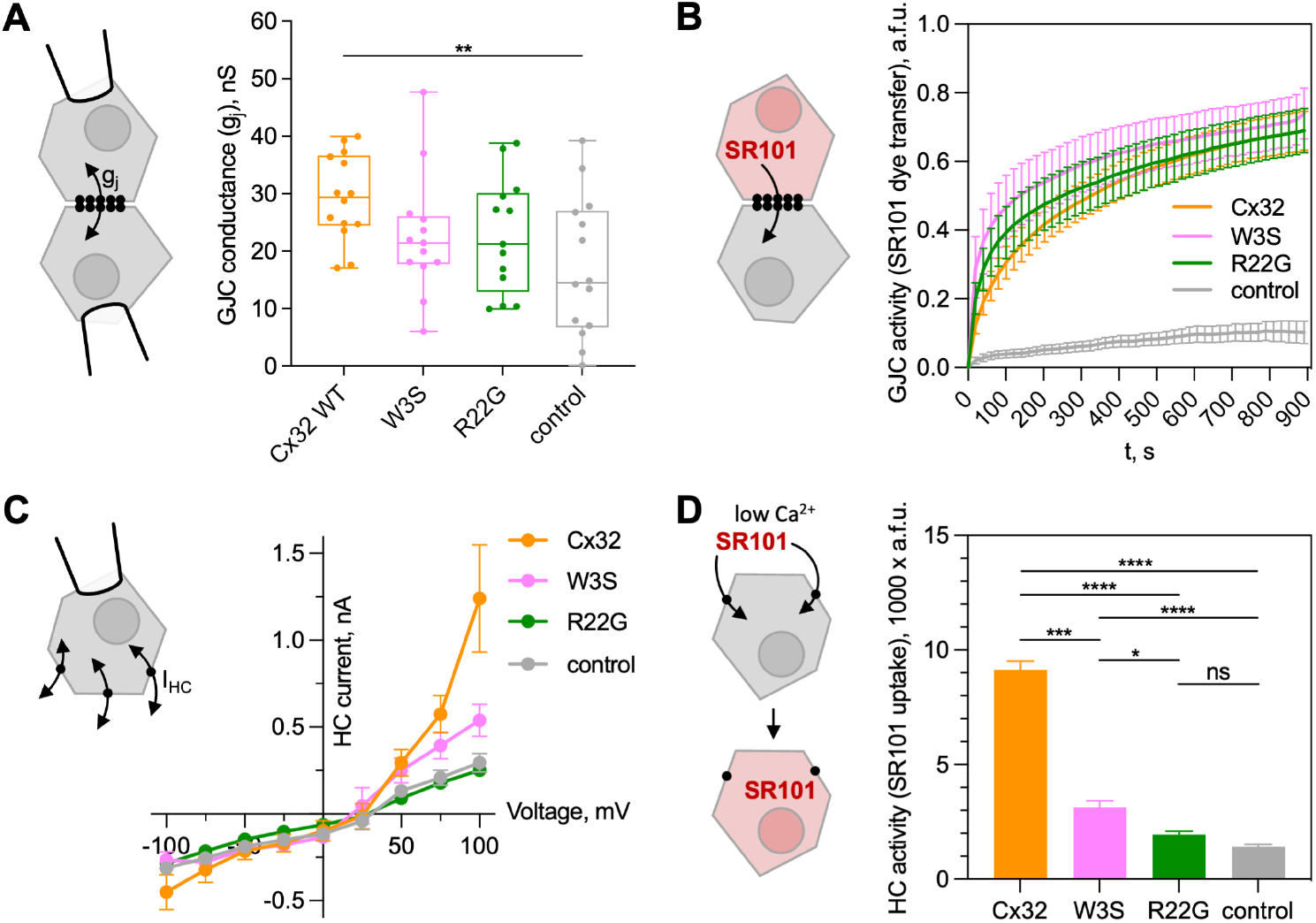
Functional properties of GJCs and HCs formed by WT, W3S or R22G Cx32 expressed in HEK293F cells. (**A**) A sketch illustrating the GJC activity measurements using dual-patch clamp; Cx32 connexons are indicated as black circles (*left*). Box plot of the GJ conductance (*g*_j_) values measured by dual-patch clamp in untransfected (control) HEK293F cells or transfected with WT, R22G or W3S constructs. HEK293F cell pairs were selected based on similar expression of the cytosolic YFP. At the end of each experiment, *g*_j_ was lowered to zero by a CO_2_-saturated extracellular solution to confirm that cells were connected by GJs and not by cytoplasmic bridges. *G*_j_ values of untransfected HEK293F cells (n = 14) resulted significantly lower (P<0.05) than WT (n = 14), but not than R22G (n = 13) and W3S (n = 13). Statistical analysis was performed using the Kruskal-Wallis test. (**B**) FRAP experiments performed in HEK293F (*left*, sketch indicating dye transfer between two coupled cells). Intercellular diffusion of the fluorescent tracer SR101 (MW 606.7 Da) was assessed in control (n = 10) and transfected cells expressing Cx32 WT (n = 20), W3S (n = 13) or R22G (n = 13). (**C**) A sketch of a whole-cell patch-clamp experiment with a cell expressing Cx32 HCs at the cell surface (*left*). Whole-cell patch-clamp experiments performed in control (n = 5) HEK293F cells or transfected with Cx32 WT (orange, n = 13), W3S (pink, n = 6) or R22G (green, n = 10). The currents mediated by mutant HCs are lower with respect to the WT. (**D**) SR101 dye uptake experiments performed in HEK293F cells not expressing (n = 935) or expressing Cx32 WT (n = 1136 cells), W3S (n = 702 cells) or R22G (n = 905 cells) HCs showed reduced uptake for the mutants. The statistical significance is indicated using asterisks (one-way ANOVA: ****, P < 0.0001; ***, P < 0.001; *, P < 0.1; ns, no significant difference).

The patch-clamp recordings were consistent with the FRAP experiments in HEK293F and HeLa DH cells (**Fig. 4B** and Fig. S17, and Fig. S16D-I, respectively). No significant difference in fluorescence recovery following 300 ms UV light exposure was observed between WT Cx32 and the W3S and R22G mutants expressed in HEK293F cells. In HeLa cells, only a mild permeability was observed in the R22G with respect to the WT, where no recovery was found in the W3S-transfected and the mock-transfected HeLa DH cells.

To assess the functional properties of the WT and mutant Cx32 HCs, we performed whole-cell patch-clamp experiments in single transfected HEK293F cells stimulated by depolarizing voltage steps (**Fig. 4C**, Fig. S18). The results highlighted a significantly lower membrane current mediated by W3S and R22G mutants with respect to the WT HCs. These observations were in line with the SR101 dye uptake experiments performed using the same cells (**Fig. 4D**, Fig. S19) and consistent with the reduced dimensions of the pore estimated by cryo-EM (**Fig. 3**).

## Discussion

The Cx32 structures put a spotlight on a debate about the conformation of the connexin NTH, which is involved in Cx32 connexon expression/assembly and in voltage gating (*20, 42, 43*). Unlike GJC structures determined previously for other connexin family members, our reconstruction of the full channel indicates that the NTH is either disordered or highly dynamic. Based on comparison with the HC structure, we hypothesize that the flexibility of the NTH in the GJC may occur due to the disruption of the NT pocket with the loss of a bound lipid molecule. The persistent presence of these six presumed sterol molecules inside the HC constricts the pore size and could limit the range of molecules that can permeate the HC. The previous work on connexin channel structure, including structure-function studies of Cx26, Cx46/50 and Cx31.1 led to a proposal of the NTH moving along the pore axis during the molecular gating events (*28, 30, 32-34*). Our structures of the WT Cx32 and W3S, R22G HCs (**Fig. 3**) show that the gating mechanism may involve an iris-like movement of the NTH (**Fig. 5**, Movie S1-2).

**Fig. 5.**
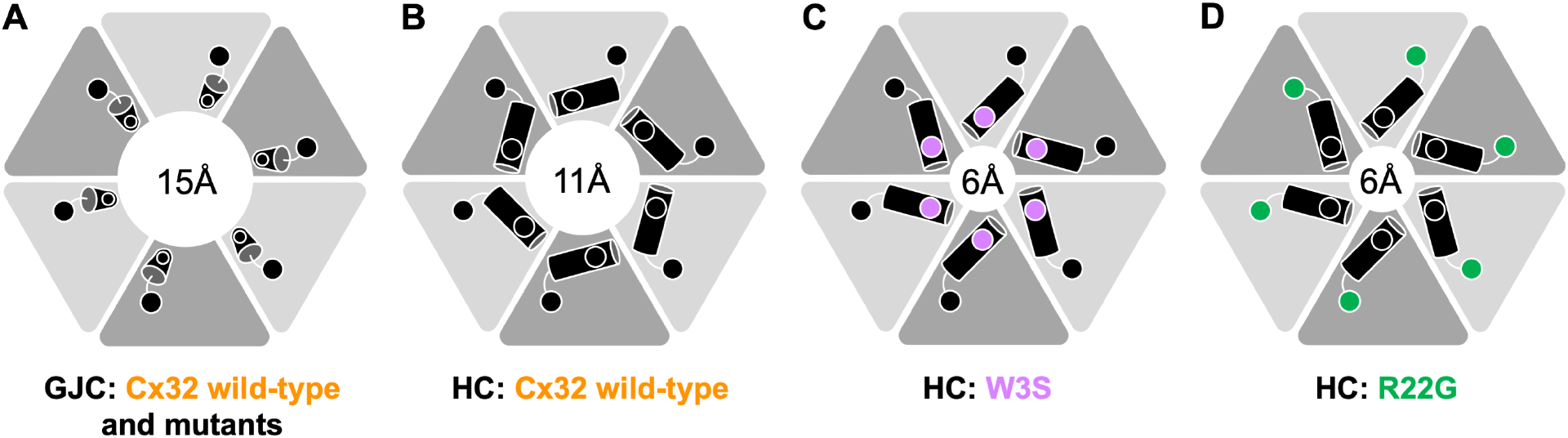
A schematic illustration of the key features observed in the Cx32 structures. (**A-D**), Cx32 GJC is determined in the open state, central pore is about 15Å. Cx32 HC is in half open state, with a central pore about 11Å.The NTH move to a conformation parallel to the membrane plane. W3S and R22G HC structures are in closed state. The NTH moves further and shrinks the pore to about 6 Å.

We propose that the flexible NTH of the GJC observed in our structure is specifically required to keep the connexon in an open state configuration to allow relatively unobstructed passage of ions, second messengers and larger diffusible molecules necessary for intercellular communication. Based on pore size considerations, the state of the WT Cx32 HC we derived may also be permeable to cytosolic ions and small molecules, supporting the notion that other cytosolic mechanisms and protein domains likely contribute to keep the Cx32 connexon in a closed state (*27, 44*). For example, the open/close transitions of the “chemical/slow” gate mediated by a cytosolic Ca^2+^ sensor have been suggested to rely on a regulator molecule, such as calmodulin, directly interacting with the Cx32 NTH and acting as a cork (*27, 45-47*).

The observation of the NTH that forms the NT pocket filled with a lipid molecule could have important implications for the gating paradigm of connexin HCs. Minute changes in the environment of the NTH (such as binding of lipids or other small molecules within the NT pocket) may lead to rearrangement of the NTH in an iris-like fashion, as seen in the Cx32 HC structure, favoring the closure of the pore at its cytosolic side. It is known that the Cx32 channel function can be modulated by hydrophobic small molecules, such as 2-APB (*48, 49*), oleamide (*50*) or anandamide (*28, 30, 32-34, 50, 51*). The possibility of manipulating the channel via alterations of the NT-pocket opens exciting opportunities for future structure-function investigations and drug screening for the treatment of CMT1X and other connexinopathies.

Our imaging and electrophysiological experiments in W3S and R22G GJCs expressed in HeLa cells confirmed previous findings that the two mutants are not functional (*37*) (*38*), despite being capable of assembling a connexon and reaching the plasma membrane. The defect can be mainly ascribed to defective formation of GJ plaques, thus to an insufficient or null number of open channels within the GJC matrix. It should be considered that most of the channels of a GJ plaque are closed under physiological conditions, especially in small plaques where it was estimated that fewer than 2% of the channels are active (*27, 52*).

Previous findings related to other CMT1X mutations (e.g., R15W (*53*) and R22P (*38*)) implicated the NTH in GJ formation. Interestingly, the expression level and functionality of W3S and R22G GJCs in HEK293F cells were similar to the WT, and only a reduced HC conductivity was observed, suggesting that the severity of the GJ defect is cell dependent (**Fig. 4**). Other Cx32 mutants, such as R75Q, M34T, V38M, R164W, Y211X, C217X were previously found to have a cell dependent expression or functionality (*20*). This underscores the need for validating the GJ activity data in more than one cell type, and ideally using more than one experimental technique.

The observation of both structural and functional alterations in the Cx32 mutant HCs expressed in HEK293F cells, but not in the GJCs, supports the recent hypothesis that Cx32 HC dysfunction may be implicated in the pathogenesis of CMT1X disease (*19, 27*). A reduced pore size in the W3S and R22G mutant HCs could conceivably impair the permeability of the channels to larger solutes such as ATP, as found in the S26L mutation (*19, 54*). Further studies are required to support the new paradigm for the molecular pathogenesis of CMT1X as being no longer linked to Cx32 GJCs but rather to Cx32 HCs. Direct observation of molecular defects in CMT1X-linked mutants is a crucial first step that sheds light on the therapeutic potential of rescuing dysfunctional Cx32 HCs as a personalized medicine strategy in the context of CMT1X disease onset and progression.

## Acknowledgments

We thank Emiliya Poghossian (EM Facility, PSI) and Miroslav Peterek (ScopeM, ETH Zurich) for expert support in cryo-EM data collection. We also thank Spencer Bliven and Marc Caubet-Serrabou (PSI) for the support in high performance computing. We also thank David Penton Ribas (Electrophysiology Facility, University of Zurich) for expert support in electrophysiology.

## Author contributions

Conceptualization: MB, VMK

Methodology: CQ, PL, EB, AV, DS, MB, VMK

Investigation: CQ, PL, EB, AV, DS, MP, VS, MB, VMK

Visualization: CQ, PL, EB, AV, DS, MB, VMK

Funding acquisition: MB, VMK

Project administration: MB, VMK

Supervision: PP, MB, VMK

Writing – original draft: CQ, PP, EV, MB, VMK

Writing – review & editing: CQ, PL, EB, AV, DS, MP, VS, PP, MB, VMK

## Data and materials availability

The coordinates and the cryo-EM density map have been deposited to Protein Data Bank and Electron Microscopy Data Bank with accession numbers: PDB ID 7ZXM, EMD-15010; PDB ID 7ZXN, EMD-15011; PDB ID 7ZXO, EMD-15012; PDB ID 7ZXP, EMD-15013; PDB ID 7ZXQ, EMD-15014; PDB ID 7ZXT, EMD-15016. The mass spectrometry data have been deposited to the ProteomeXchange Consortium via the PRIDE partner repository with the dataset identifiers PXD039379, PXD033848 and PXD033671.

## Supplementary Materials

Materials and Methods

Figs. S1 to S19

Table S1

References (*55*–*75*)

Movies S1 to S2

