## Supplementary Information for "Structures of wild-type and selected CMT1X mutant connexin 32 gap junction channels and hemichannels"

### Materials and Methods

**Molecular cloning and mutagenesis.** Human connexin 32 (Cx32, Uniprot ID P08034), followed by a C-terminal YFP and twinStrep tag, was cloned into a pACMV plasmid. The W3S and R22G mutations were introduced using site-directed mutagenesis. For IRES (internal ribosomal entry site) containing constructs, wild-type untagged Cx32 and its mutants W3S and R22G were subcloned using NheI and MreI into MCS site prior to the IRES sequence followed by YFP sequence containing pACMV vector.

#### Cell culture

Large-scale protein purification: Human Embryonic Kidney Freestyle 293F (HEK293F; Thermo Fischer Scientific) cells were cultured in 15 cm cell culture dishes containing Dulbecco's Modified Eagle's medium (DMEM, BioConcept) supplemented with 10% FBS and 1% penicillin/streptomycin (PenStrep, PAN Biotech), 5% $CO_2$ and cultured at 37 °C. At 80% confluence, cell medium was exchanged to DMEM supplemented with 2% FBS and 1% PenStrep. The transfection mixture was prepared with Cx32-YFP-twinStrep plasmid and branched polyethyleneimine (PEI, Sigma-Aldrich) at a ratio of 1:2 (w/w) with serum-free DMEM. The transfection mixture was added to cells after 5 min of incubation at room temperature. After 48 h of expression at 37 °C and 5% $CO_2$, the cells were harvested using a cell scraper in buffer A containing 25 mM Tris-HCl, pH 8.0, 150 mM NaCl) and collected by centrifugation at 800 g and 4 °C. The cells were washed with buffer A and collected by centrifugation. The supernatant was discarded, and the cells frozen at -80 °C for further experiments.

Surface biotinylation: HEK293F cells were cultured and transiently transfected with IRES construct as described for large-scale expression. HeLa DH cells were cultured like HEK293F cells and transiently transfected with 45 μg of IRES construct plasmid per plate using Lipofectamine 3000 (Thermo Fischer Scientific) according to manufacturer's protocol.

Immunocytochemistry and western blot: HEK293F or HeLa DH (DH clone, Sigma-Aldrich) cells were seeded on poly-L-lysine (Cultrex) precoated glass coverslips in 6-well plates at a density of 0.7 million cells per well. For western blot (WB) and whole-cell patch-clamp experiments HEK293F cells were cultured as in ICC without the glass slide. The cell count was determined using trypan blue viability staining and the cells seeded in DMEM supplemented with 10% FBS and 1% PenStrep and cultured at 37 °C and 5% $CO_2$. 24 h after seeding, HEK293F cells were transiently transfected as described above using 4 μg DNA per well. HeLa DH cells were transiently transfected with 3 μg DNA using Lipofectamine 3000 according to the manufacturer's protocol. The cells were placed at 37 °C and 5% $CO_2$ for 36 h of expression.

Dye transfer assays: For dye transfer assays in HEK293F cells, the cells were seeded and cultured as described above at 30 000 cells per well in ibidi 8-well μ-Slide chambers or 300 000 cells per ibidi μ-Dish, precoated with poly-L-lysine. 24 h after seeding, HEK293F cells were transfected as described above using 0.46 μg of DNA per well of a chamber and 2 μg of DNA per dish. The cells were placed at 37 °C and 5% $CO_2$ for 36 h of expression. HeLa DH cells were plated onto 12 mm diameter glass coverslips. 24 h after plating, the cells were transiently transfected with 0.2 µg of the DNA using IRES constructs using Lipofectamine 3000 according to manufacturer's protocol and were placed at 37 °C and 5% $CO_2$ for 24 h of expression.

Electrophysiology assays in HeLa cells: A clone of HeLa cells (HeLa DH) that is essentially devoid of connexins was utilized in our experiments. HeLa DH cells were cultured in DMEM/F12 medium (Gibco) supplemented with 10% FBS (Gibco) and 1% PenStrep at 37 °C in a humidified incubator with 5% $CO_2$. 24 h after plating cells onto 12 mm diameter glass coverslips, the cells were transiently transfected with 0.2 µg of the DNA of Cx32-IRES-YFP (WT) or mutant vectors (R22G or W3S) using Lipofectamine 3000 (Thermo Fisher Scientific). The mixture was added to the wells drop by drop and left to act for 4 hr, after which, the medium was replaced. The mixture applied to control (*Mock*) cells did not contain DNA. The experiments took place 24 h post transfection.

Electrophysiology assays in HEK293F cells: HEK 293F cells were cultured in DMEM medium (Gibco) supplemented with 10% FBS (Gibco) and 1% PenStrep at 37 °C in a humidified incubator with 5% CO2. The cells were seeded onto a 24–well dish and incubated for 18-24 hours at 37 °C until they were approximately 50-70% confluent. After which, the cells were transiently transfected with 0.2 µg of the DNA of Cx32-IRES-YFP (WT) or mutant vectors (R22G or W3S) using Rotifect (Carl Roth) according to standard manufacturer's protocol. 24 h after the transfection, the cells were seeded on glass coverslips (previously coated with polyornithine 100µg/ml for 1 hr at RT). The experiments took place 48 h post transfection.

**Cx32 purification.** The cells were resuspended in Buffer A supplemented with protease inhibitors (1 mM benzamidine, 1 µg/ml leupeptin, 1 µg/ml aprotinin, 1 µg/ml pepstatin, 1 µg/ml trypsin inhibitor and 1 mM PMSF). Cells were lysed using sonication at 35% amplitude with two sonication cycles per plate (cycle: 0.5 s pulse on, 0.5 s pulse off), and the membrane preparation was clarified by ultracentrifugation at 35000 rpm (Beckman Ti45 rotor) to remove the soluble fraction. The membrane pellet was resuspended in Buffer A supplemented with protease inhibitors (1 mM benzamidine, 1 µg/ml leupeptin, 1 µg/ml aprotinin, 1 µg/ml pepstatin, 1 µg/ml trypsin inhibitor and 1 mM PMSF) using a homogenizer. The membranes were solubilized by adding 1% DDM and 0.2% cholesteryl hemisuccinate (CHS) and incubated at 4 °C for 1 h. The solubilized membranes were centrifuged by an additional round of ultracentrifugation. The supernatant was incubated with 1 ml of CNBr-activated Sepharose coupled anti-GFP nanobody (*55*) at 4 °C. The resin and protein mixture was collected in a gravity column and washed with 40 column volumes of Buffer B (25 mM Tris-HCl, pH 8.0, 150 mM NaCl, 0.1% digitonin). Protein was cleaved overnight at 4 °C by adding 0.4 mg HRV 3C protease. The Cx32 protein was collected and concentrated to 1 ml using a 100 kDa cut-off concentrator. The Cx32 sample was injected onto a Superose 6 Increase 10/300 GL column pre-equilibrated with Buffer B. The Cx32 peak fractions were collected, concentrated, and used for cryo-EM grid preparation immediately. The expression and purification methods of W3S and R22G mutants were the same as the wild type.

**Cryo-EM sample preparation and data collection.** The purified Cx32 wild-type protein, W3S, and R22G were concentrated to about 5 mg/ml. The cryo-EM grids were glow-discharged for 30 s using an PELCO easiGlow™ glow discharge system. 3.5 μl of the protein was added to the glow-discharged Quantifoil R1.2/1.3 200-mesh grid. The grids were blotted using Vitrobot Mark IV (Thermo Fisher Scientific) and plunge-frozen in liquid ethane (blot force 10, blot time 3 s). The grids were transferred to liquid nitrogen for storage and cryo-EM data collection. Cryo-EM datasets were collected using a Titan Krios electron microscope equipped with a K3 direct electron detector (Gatan) and a GIF-Quantum energy filter (slit width of 20 eV) at ScopeM, ETH Zurich. The defocus range was set from -1.0 μm to -2.0 μm. The micrographs were collected using EPU2.0 software and dose-fractioned to 40 frames in super-resolution mode. The exposure time for each micrograph was 0.7 s, and the total dose was about 50 $e^-/Å^2$.

**Cryo-EM data processing.** The images were classified optics groups using a script provided by Dr. Pavel Afanasyev (ETH Zurich; https://github.com/afanasyevp/afis). The movies were first binned two-fold and motion-corrected using MotionCor2 (*56*). The nominal pixel size of the micrograph after two-fold binning was 0.678 Å. Gctf (*57*) was used for CTF estimation. About 1000 particles were manually picked in Relion3.1 (*58, 59*). 2D classification of the manually picked particles showed clear features of gap junction channels (GJCs) and hemi channels (HCs). The selected 2D classes were used as templates to autopick the particles in all micrographs using Relion3.1. After several rounds of 2D classification, the GJCs and HCs were selected separately based on their feather and transferred to Cryosparc2 (*60*) to generate the initial models. W3S and R22G were processed using Cx32 as initial model. 3D classification of GJCs and HCs was performed separately in Relion3.1. The good 3D classes for GJCs and HCs were selected for further 3D refinement. CTF refinement and particle polishing were also performed using Relion3.1 to further improve the resolution. The calibrated pixel size was corrected to 0.654 Å during postprocessing. Local resolution maps were calculated using ResMap (*61*) implemented in Relion 3.1. The detailed steps of image processing are shown in Fig. S4-7 and Table S1.

**Model building and refinement.** Model building was carried out manually in COOT (*62*). Connexin26 structure (PDB ID: 2ZW3) was used as a guide template and the side chains of the bulky residues (Phe, Trp, Tyr and Arg) were used as a guide for model building. The model was refined using real_space_refine in PHENIX (*63*). The model was validated as previously described (*64*). The atoms of the refined model were randomly displaced by 0.5 Å using PDB tools implemented in PHENIX. This perturbed model was refined against the half map1. The new refined model was used to generate FSC curve of model versus half map2. The geometries of the model were validated using MolProbity (*65*). All figures were prepared in PyMol (*66*), Chimera (*67*) and Chimera X (*68*).

**Mass spectrometric analysis of purified Cx32 protein N-terminus**

<u>*Sample preparation for LC-MS/MS analysis:*</u> 25 μg of protein (Cx32, W3S and R22G) were digested using ProtiFi S-Trap™ micro columns according to the manufacturer's instructions. The peptides were dried in a vacuum centrifuge and resuspended in 5% I, 0.1% FA.

<u>*LC-MS/MS data acquisition:*</u> The sample was injected and analyzed on an Orbitrap Exploris™ 480 mass spectrometer, equipped with a nanoelectrospray source and coupled to an Easy-nLC 1200 (Thermo Fisher Scientific). Peptides were separated with a 60 min gradient from 3-30% B (Eluent A: 0.1% FA, Eluent B: 95% ACN, 0.1% FA) on a 40 cm x 0.75 μm in-house packed C18 column (see above for details), heated to 50 °C. The three samples were measured in data

independent acquisition (DIA) mode with 41 fragmentation windows with 1 m/z overlap. MS1 scan range was set to 350-1150 m/z with a normalized AGC target of 200% and RF Lens of 50%. The maximum injection time was set to 264 ms and the Orbitrap resolution to 120,000. For MS2 scans the normalized AGC target was set to 200%, the maximum injection time was 66 ms, RF Lens 50% and the collision energy was set to 30%. The total method length was 72 min.

Data analysis: The acquired raw files were analyzed with Spectronaut™ version 15.7 in directDIA™ mode. Default settings were used, except that iRT calibration, cross run normalization and imputation were switched off and single hits were excluded. For identification a FASTA file of the human proteome (downloaded from UniProt (*69*)), as well as a FASTA file with all Cx32 sequences (mutations and WT), as well as a contaminant FASTA file (MaxQuant (*70*)) were used. Plots were made in R v. 4.1.1 using the R packages tidyverse (*71*) and protti (*72*). Raw files, file annotations, FASTA files and search results have been deposited on PRIDE (*73*): PXD039379 (surface biotinylation, HeLa DH), PXD033848 (purified protein analysis), PXD033671 (surface biotinylation, HEK293F).

**Surface protein biotinylation.** Cell surface protein biotinylation was done using Pierce™ Cell Surface Biotinylation and Isolation kit (Thermo Fischer Scientific) according to manufacturer's protocol. The eluted surface biotinylated proteins were analyzed using mass spectrometry.

**Mass Spectrometric analysis of surface protein biotinylation, HEK293F cells**

*Sample preparation for LC-MS/MS analysis*: 100 µL of each eluted sample were digested using ProtiFi S-Trap™ micro columns with a few modifications of the manufacturer's protocol. Briefly, disulfide bonds did not have to be reduced with dithiothreitol (DTT) as the elution buffer already contained DTT. 50 µL of 2x lysis buffer (10% SDS, 100 mM TEAB pH 7.55) were added and free cysteines were alkylated with 40 mM iodoacetamide. The rest of the sample preparation was conducted as recommended by the manufacturer. After drying, the peptides were resuspended in 30 µL 5% ACN, 0.1% FA with an addition of iRT peptides (Biognosys). For library generation, pooled samples of each condition were measured in DDA and included alongside all DIA measurements of all samples.

*LC-MS/MS data acquisition, DDA measurements:* Peptides from pooled samples of each condition were separated with a linear gradient from 3 to 35% B (eluent A: 0.1% FA in water, eluent B: 0.1% FA IACN) on an in-house packed 40 cm x 0.75 µm C18 column (1.9 µm C18 beads, Dr. Maisch Reprosil-Pur) connected to an Acquity UPLC M-Class system (Waters). Samples were acquired on a Thermo Orbitrap Fusion™ Lumos™ Tribrid™ mass spectrometer at a normalized AGC target of 200% for both MS1 and MS2. The scan range was set to 350-1150 m/z, RF Lens to 30%, the maximum injection time was 100 ms for MS1 and 54 ms for MS2. Cycle time was 3 s and dynamic exclusion was set to 60 s. Charge states between 2 and 7 were acquired, the Orbitrap resolution for MS1 was set to 120,000 and 30,000 for MS2. Peptides were fragmented with HCD at a collision energy of 30%. The total method length was 165 min.

*LC-MS/MS data acquisition, DIA measurements:* Peptides were separated on a similar gradient from 3-35% B and a total method length of 165 min. MS1 scan range was set to 250-1400 m/z and RF Lens to 30%. Normalized AGC target was 50% and the maximum injection time for MS1 was 100 ms. MS1 Orbitrap resolution was 120,000, MS2 Orbitrap resolution was set to

30,000. Peptides were fragmented in 41 sequential windows with 1 m/z overlap, with a maximum injection time of 54 ms. HCD collision energy was set to 28%.

Data analysis: Spectronaut v. 15.7 was used for data analysis. Libraries were generated from all samples measured in DDA and DIA. CAMthiopropanoylation (+ 145.02 Da) was included as a PTM, Trypsin/P and Lys-C/P were set as digestion enzymes. The samples were searched against sequences of the human proteome (downloaded from UniProt (*69*)), mutated and non-mutated Cx32, CFP and a FASTA file of common contaminants (MaxQuant (*70*)). Peptides identified in mutated regions were excluded from the quantification. The minimum peptide length was set to 5 amino acids. During the search, single hits were excluded and PTM localization was conducted. Missing values were not imputed. Data was exported from Spectronaut and further analyzed in R v. 4.1.1 using the R packages tidyverse (*71*), protti (*72*) and ggpubr (*74*). Raw files, file annotations, FASTA files, spectral library, search results and the data analysis script are deposited on PRIDE (PXD033671).

**Mass Spectrometric analysis of surface protein biotinylation, HeLa cells**

*Sample preparation for LC-MS/MS analysis*: 50 µL of each sample were digested with ProtiFi S- Trap™ micro columns as mentioned above. Pooled samples of each condition were prepared for DDA library measurements and non-biotinylated control samples were included to exclude proteins that could show unspecific binding.

*LC-MS/MS data acquisition, DDA measurements*. Peptides from pooled samples of each condition were separated with a linear gradient from 3 to 30% B (eluent A: 0.1% FA in water, eluent B: 95% ACN, 0.1% FA) on an in-house packed 40 cm x 0.75 µm C18 column (3 µm C18 beads, Dr. Maisch ProntoSIL) connected to an easy-nLC 1200 (Thermo Fisher Scientific). Measurements were acquired on a Thermo Orbitrap Exploris™ 480 mass spectrometer at a normalized AGC target of 200% for both MS1 and MS2. The scan range was set to 350-1150 m/z, RF Lens to 50%, the maximum injection time was 264 ms for MS1 and 54 ms for MS2. 20 dependent scans were acquired and ynamic exclusion was set to 20 s. Charge states between 2 and 6 were selected, the Orbitrap resolution for MS1 was set to 120,000 and 30,000 for MS2. Peptides were fragmented with HCD at a collision energy of 30%. The total method length was 132 min.

*LC-MS/MS data acquisition, DIA measurements.* All samples (biotinylated samples and non-biotinylated controls) were measured in DIA. Peptides were separated on a similar gradient from 3-30% B and a total method length of 132 min. MS1 scan range was set to 350-1150 m/z and RF Lens to 50%. Normalized AGC target was 200% and the maximum injection time for MS1 was 264 ms. MS1 Orbitrap resolution was 120,000, MS2 Orbitrap resolution was set to 30,000. Peptides were fragmented in 41 sequential windows with 1 m/z overlap, with a maximum injection time of 66 ms. HCD collision energy was set to 30%. A lock-mass (445.12003 m/z) was included for internal mass calibration.

Data analysis: Data was analyzed as mentioned above, except that non-biotinylated samples were searched and the identified proteins (unspecific binders) were excluded from the analysis.

**Western blot analysis (WB).** Cells, analyzed for total protein expression, were washed with 1X PBS, trypsinized with 500 µl of trypsin-EDTA (BioConcept) for 5 min, resuspended in 500 µl of ice-cold DMEM supplemented with 10% FBS and 1% PenStrep and transferred to a microcentrifuge tube. The cells were pelleted by centrifugation for 20 min at 1000 rpm at 4 °C and the supernatant was discarded. The pellet was resuspended in 150 mM NaCl, 50 mM Tris-

HCl pH 8.0 supplemented with DNase I. The cells were sonicated for five cycles as described in protein purification and the cells were diluted in 4X SDS-PAGE loading buffer.

**Immunocytochemistry (ICC).** The cells were fixed on a glass coverslip with 4% formaldehyde in 1X PBS at room temperature for 20 min and washed twice with 1X PBS. In experiments evaluating protein localization, the plasma membrane was additionally stained with Vybrant CM-DiI (Invitrogen) for 10 min, washed three times with 1X PBS and fixed again as described above. The coverslips were transferred to buffer containing 5% urea [w/v], 100 mM Tris-HCl pH 8.0 at 95 °C for 10 min for antigen retrieval. The coverslips were washed three times with 1X PBS for 5 min and then placed in 1X PBS containing 100 μM digitonin for 10 min for membrane permeabilization, followed by the same washing procedure as described above. The coverslips were then incubated in 1% BSA, 22.52 mg/ml glycine in PBST (0.1% Tween 20) buffer for 30 min and then transferred to 1% BSA in PBST buffer containing rabbit polyclonal anti-Cx32 antibody (#34-5700, Invitrogen) at a 1:250 dilution in a humidified chamber at 4 °C for overnight incubation. The coverslips were again washed three times by 5 min incubation in 1X PBS and transferred to 1% BSA PBST buffer containing goat anti-rabbit IgG Alexa Fluor® 488-conjugated secondary antibody (ab150077, Abcam) at 1:1000 dilution for protein localization assessment, or goat anti-rabbit Alexa Fluor® 647-conjugated secondary antibody (ab150079, Abcam) at 1:2000 dilution for connexin and YFP co-expression assessment. The cells were then washed three times with PBS for 5 min, stained with 50 μg/ml Hoechst33342 fluorescent dye (Sigma) in 1X PBS for 1 min, washed once with 1X PBS and then mounted on glass slides using gelvatol. The samples were stored at 4 °C prior to imaging.

**Dye transfer assays.** For dye uptake assays in HEK293F cells, the cells were washed twice with PBS-E (1X PBS 5 mM EDTA) and pre-incubated with PBS-E for 5 min. The PBS-E was then replaced with PBS-E containing 50 μg/ml sulforhodamine 101 (SR101, Sigma) and incubated 5 min for HC dye-uptake assay and 15 min for GJC FRAP assay. After incubation, the cells were washed four times with 1X PBS and imaged in FluoroBrite™ DMEM supplemented with 1% FBS and 1% PenStrep.

For dye uptake experiments in HeLa DH cells, the cells were loaded with 10 μM of Calcein Blue AM (Invitrogen) dye in DMEM/F12 medium in the presence of 25 μM sulfinpyrazone (Sigma-Aldrich) and 0.1% of Pluronic F-127 (Invitrogen) and incubated at 37 °C in a humidified incubator with 5% $CO_2$ for 30 mins, before imaging in an extracellular solution containing 150 mM NaCl, 10 mM HEPES, 5 mM KCl, 5 mM glucose, 1 mM $MgCl_2$, 2 mM $CaCl_2$ and 2 mM sodium pyruvate (pH 7.4, 311 mOsm).

**Light microscopy.** ICC and HEK293F dye transfer assay samples were imaged using Leica STELLARIS 5 confocal microscope using LAS X (4.2.1.23819 – build 23180) acquisition software. The overview images, where applicable, were acquired using HC PL APO 20X/0.75 dry objective and the images for data analysis with a HC PL APO 63X/1.4 oil CS2 objective and a pinhole airy of 1.06 AU (101.4 μm), pixel dwell time of 3.1625 μs and a scan speed of 400 Hz. The images were collected using the 405 nm, 488 nm, 561 nm and 638 nm laser lines laser lines for Hoechst 33342, EYFP and Alexa488, SR101 and CM-DiI, and Alexa647 fluorescent dyes respectively with a frame sequential data acquisition scheme and detected with a HyD camera detector.

ICC samples: For GJ characterization, overview images of the whole coverslip were collected with 10% overlap and stitched in the acquisition software. The positions for data collection were distributed evenly over the whole surface where respective connexin and plasma membrane signals were present to avoid bias as well as target transfected cells. Z-stacks of 7 µm were collected with a z step of 0.3 µm from the defined positions, with the center of each stack determined as the z height with the highest signal from the nucleus. For non-quantitative ICC samples, the data was collected without prior overview acquisition.

*HC dye uptake assays in HEK293F cells:* 10 images were collected per condition and each condition was imaged using same microscope image acquisition settings. Each assay was done in experimental triplicates.

*GJC FRAP assays in HEK293F cells:* The YFP fluorescence was used to identify transfected cells. Neighboring cells with YFP fluorescence were selected as regions for FRAP experiment. A cell bordering to other cells is defined as the ROI and bleached with 100% SR101 laser line intensity after collecting six pre-bleach images at a lower SR101 laser line intensity in a 5.16 s intervals. After bleach pulses, the fluorescence recovery was recorded at the lower SR101 laser line intensity every 20.16 s for 45 frames, where the dye recovery reached plateau.

*GJC FRAP assays in HeLa DH cells:* The cells were mounted on the stage of an upright wide-field fluorescence microscope (Olympus BX51WI) with an infinity-corrected water immersion objective (40X, 0.8 NA, Olympus). Cytosolic YFP fluorescence was excited by a 470 nm LED while Calcein Blue fluorescence was excited by a 365 nm LED. Isolated pairs of Cx32 transfected HeLa cells were selected based on their cytosolic YFP expression. A 375 nm UV laser beam was focalized for 300 ms on one cell of the pair to bleach its Calcein Blue content and observe the following recovery mediated by passive diffusion of Calcein Blue through Cx32 GJs connecting the cell pair.

**Light microscopy image analysis**

*GJ characterization:* Stacks collected from the same region but with different channels were imported in Fiji and merged. Connexin signal was considered as a GJ if it colocalized with the plasma membrane stain and was present in three or more stack slices. The GJs were counted and measured from the longest slice manually. The measurements were exported to GraphPad Prism and compared using one-way ANOVA, followed by Tukey's multiple comparisons test.

*HC dye transfer assays in HEK293F cells:* Images were processed using a python script (https://github.com/lavrihap/hc-data-processing.git). The values were imported to GraphPad Prism 9.0.0 for analysis. The replicates were combined and plotted as mean ± SEM. The significance of dye uptake difference between different conditions was assessed using ordinary one-way ANOVA and Games-Howell's multiple comparisons test.

*GJC FRAP dye transfer assays in HEK293F cells:* FRAP images were imported in Fiji and pixel intensities in the time series were measured from three regions – bleached cell (ROI1), neighboring cells including the bleached cell (ROI2) and background (ROI3). The fluorescence recovery curves were calculated with EasyFRAP-web software (*75*) using full-scale normalization and the normalized curves exported to GraphPad Prism. All curves were plotted, and a one-way association curve was fitted to the data. The percentage of FRAP recovery was determined and compared using ordinary one-way ANOVA and Tukey's multiple comparisons test with a single pooled variance.

*GJC FRAP dye transfer assays in HeLa DH cells:* Data were analyzed using a software we developed in Matlab (The MathWorks, Inc.).

**Electrophysiology**

Dual patch-clamp electrophysiological experiments in HeLa DH and HEK293F cells: A glass coverslip of Cx32-IRES-YFP (WT) or mutant (R22G or W3S) transfected cells was transferred to an experimental chamber at room temperature (22–24 °C) and mounted on the stage of an upright wide-field fluorescence microscope (Olympus BX51WI) with an infinity-corrected water immersion objective (40X, 0.8 NA, Olympus). We continuously superperfused the cells at the rate of 2 ml/min with an extracellular solution containing 130 mM NaCl, 10 mM HEPES, 5 mM KCl, 5 mM glucose, 1 mM $MgCl_2$, 2 mM $CaCl_2$, 2 mM sodium pyruvate, 4 mM TEA-Cl, 4 mM CsCl, 2 mM 4-Aminopyridine, 2 µM TRAM and 400 nM UCL (pH 7.4, 318 mOsm). Cytosolic YFP fluorescence was excited by a 470 nm LED to identify the transfected cells and allow subsequent electrophysiological analysis. Patch-clamp recordings were performed using an Axon 700B amplifier (Molecular Device) with a dual headstage configuration capable of carrying out simultaneous measurements. Pipettes were filled with an intracellular solution containing 125 mM KAsp, 10 mM NaCl, 10 mM HEPES, 13 mM KCl, 1 mM $MgCl_2$ and 50 µM BAPTA (pH 7.2, 309 mOsm) filtered through a 0.22 µm pore size membrane (Millipore). The resistance of patch pipettes in the bath ranged between 6 to 8 MOhm. Once the whole-cell configuration was achieved, both cells were voltage-clamped at -20 mV. The junctional current $I_j$ was measured in cell 1 by applying voltage steps $V_j = +10$ mV to cell 2. The corresponding junctional conductance was computed as $g_j = I_j/V_j$. At the end of almost all experiments, we applied $CO_2$ to the bath to prove that junctional currents occurred through Cx32 GJ WT or mutant channels. Electrophysiological data were acquired by pClamp software (version 10.4, Molecular Device) and analyzed with a software we developed in Matlab (The MathWorks, Inc.).

Whole-cell patch clamp experiment: HC activity was measured using whole-cell patch-clamp with EPC10 amplifier (HEKA, Germany). Data were acquired at 1 ms sampling rate and filtered at 3KHz. Isolated cells expressing Cx32, W3S and R22G were patched. Cells were continuously perfused with extracellular solution containing 140 mM NaCl, 5.4 mM CsCl, 1.8 mM $CaCl_2$, 1 mM $MgCl_2$, 2 mM $BaCl_2$, 10 mM HEPES (~310 mOsm) at pH 7.4 adjusted with NaOH. The 5.4 mM CsCl and 2 mM $BaCl_2$ was added to suppress $Na^+$-$K^+$-ATPase pump activities and $Ca^{2+}$-activated $K^+$ channel currents. Patch pipettes were pulled from borosilicate glass at a resistance of 3-4 MΩ and was filled with intracellular solution containing 130 mM CsCl, 10 mM Na Gluconate, 0.26 mM $CaCl_2$, 1 mM $MgCl_2$, 2 mM EGTA, 7 mM TEACl and 5 mM HEPES (~300 mOsm) at pH 7.2 adjusted with CsOH. The 130 mM CsCl was added, where $Cs^+$ was sole charge carrier that do not mediate outward current through $K^+$ channels and TEA.Cl was used to block residual $K^+$-currents. All recordings were performed at room temperature. After 45-60 s of whole-cell giga seal configuration the HC activity was measured with voltage steps $\Delta V = 25$ mV from -100 to +100 mV was applied for every 1 s for 10 s. Data was analyzed using fit master (HEKA, Germany) and peak amplitudes were measured to generate I-V curve. Statistical analysis on peak amplitude current at different voltage steps were performed using ordinary one-way ANOVA and Sidak's test. Data are presented as mean ± SEM.

### Supplementary Figures

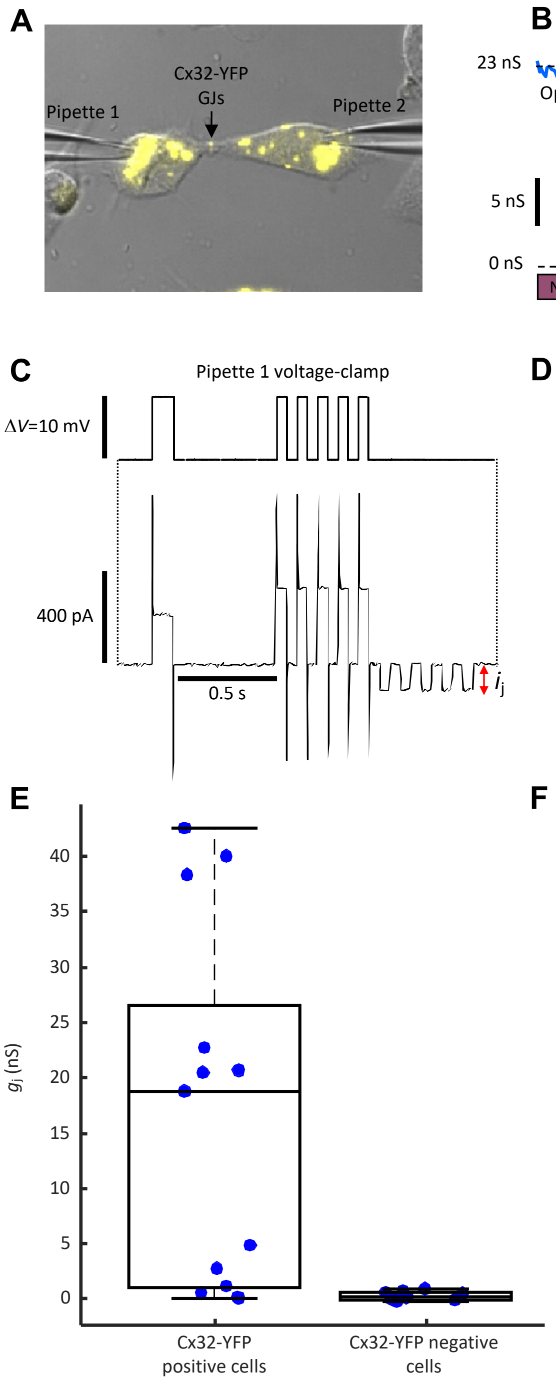


**Figure S1. Monitoring Cx32-YFP GJC conductance ($g_j$).** (**A**) Dual patch clamp electrophysiology to measure the $g_j$ in HeLa cell pairs transfected with the Cx32-YFP plasmid. The arrow indicates Cx32 GJC between two HeLa cells. (**B**) $g_j$ was measured while perfusing cells with normal extracellular solution (NS) or NS saturated with $CO_2$ to verify that the junctional current $i_j$ was mediated by GJC. (**C-D**) Voltage patch clamp protocols: both cells were stepped by $\Delta V = 10$ mV from the –20 mV holding potential to measure the cell parameters and $i_j$. The junctional resistance was computed as $R_j = \Delta V / i_j$, where $\Delta V$ was corrected by the voltage drop due to pipette resistance. The junctional conductance was then measured as $g_j$ = 1/ $R_j$. (**E**) Box plot of $g_j$ measured in YFP-positive (n = 13) and YFP negative (n = 9) HeLa cell pairs with mean ± STD = 16.4 ± 16.2 nS vs. 0.3 ± 0.4 nS, respectively. Statistical analysis was performed using the Mann-Whitney *U* test (P = 0.002). (**F**) Time course of the junctional conductance $g_j$ (n = 13 cell pairs) and membrane conductance $g_m = 1/R_m$ (n = 25 cells) during perfusion with NS saturated with $CO_2$ followed by NS.

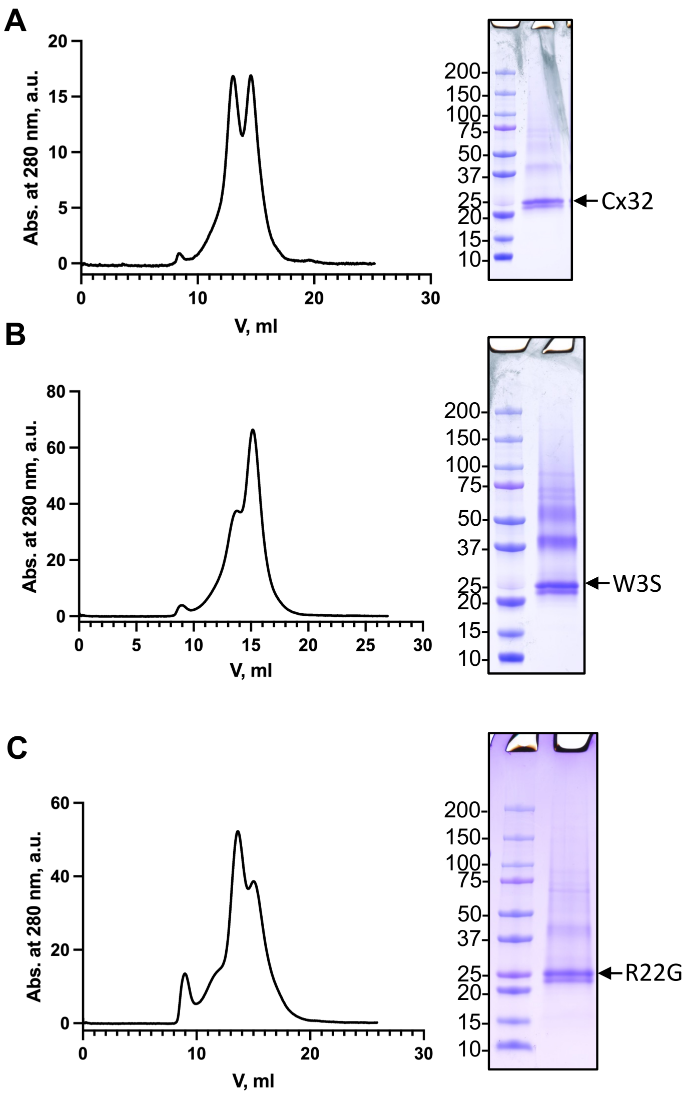


**Figure S2. Expression and purificaiton of Cx32 and mutants.** (**A**) Size-exclusion chromatogram (SEC) and Coomassie stained SDS-PAGE gel of Cx32 purification. (**B-C**) SEC and SDS-PAGE gel of the W3S and R22G samples purified with the same protocol as Cx32 wild type.

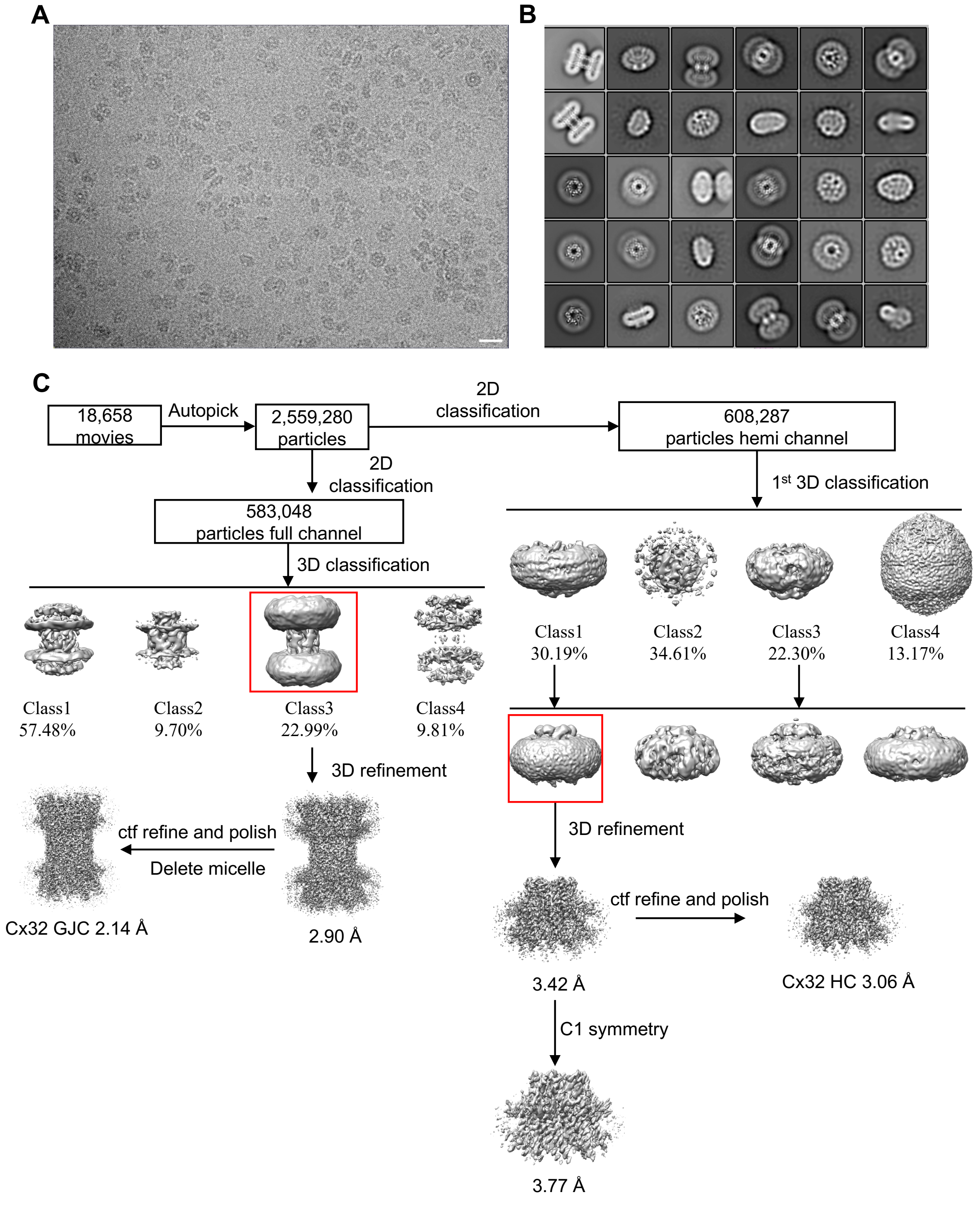


**Figure S3. Cryo-EM image processing pipeline of Cx32 dataset.** (**A**) A representative micrograph of Cx32 cryo-EM sample. Scale bar = 200 Å. (**B**) Images of reference-free 2D classification of Cx32. The box size is 251 Å. (**C**) Overview of the Cx32 GJC and HC data processing pipeline.

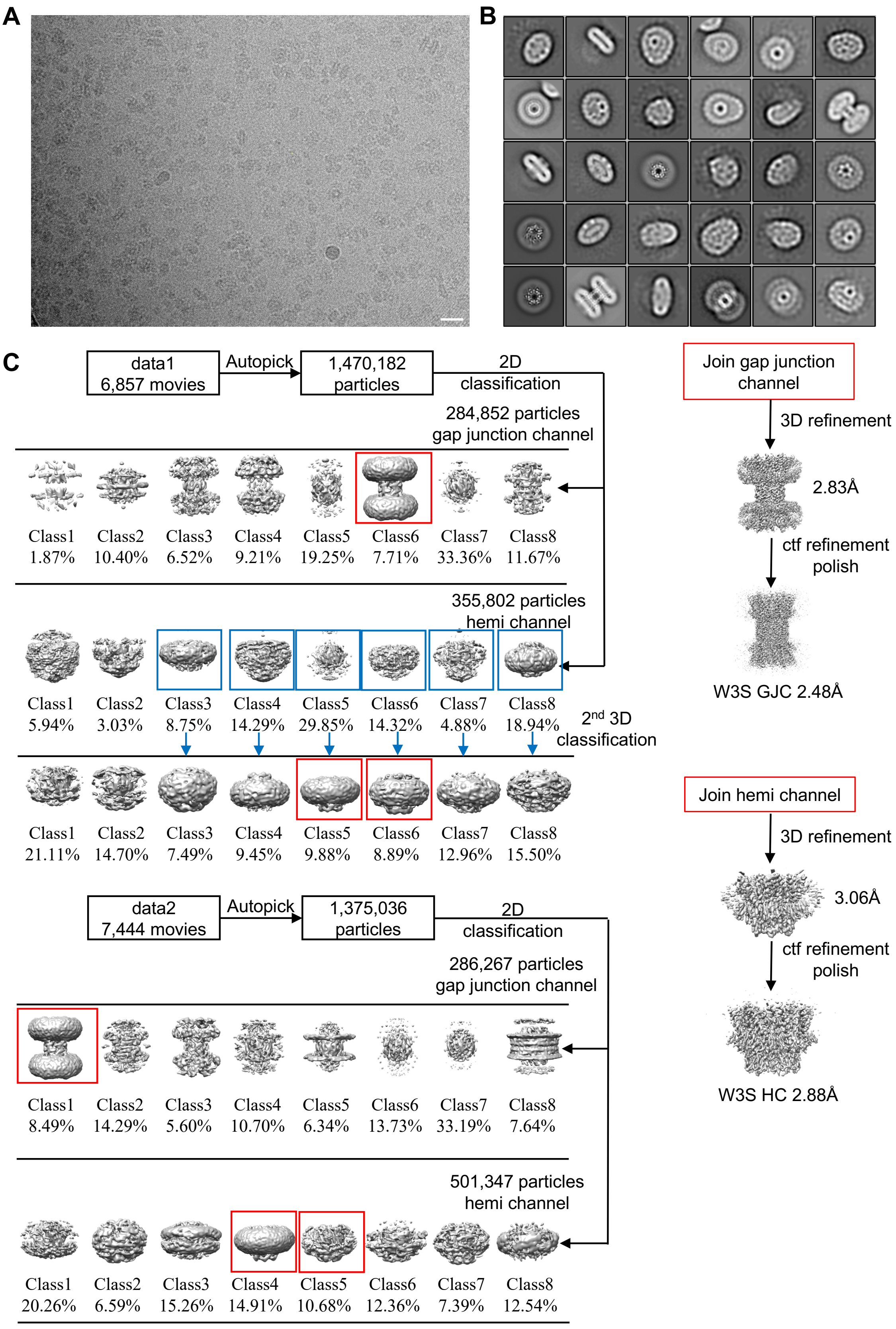


**Figure S4. Cryo-EM image processing pipeline of W3S dataset.** (**A**) A representative micrograph of W3S cryo-EM sample. Scale bar = 200 Å. (**B**) Images of reference-free 2D classification of W3S. The box size is 251 Å. (**C**) Overview of the W3S GJC and HC data processing pipeline.

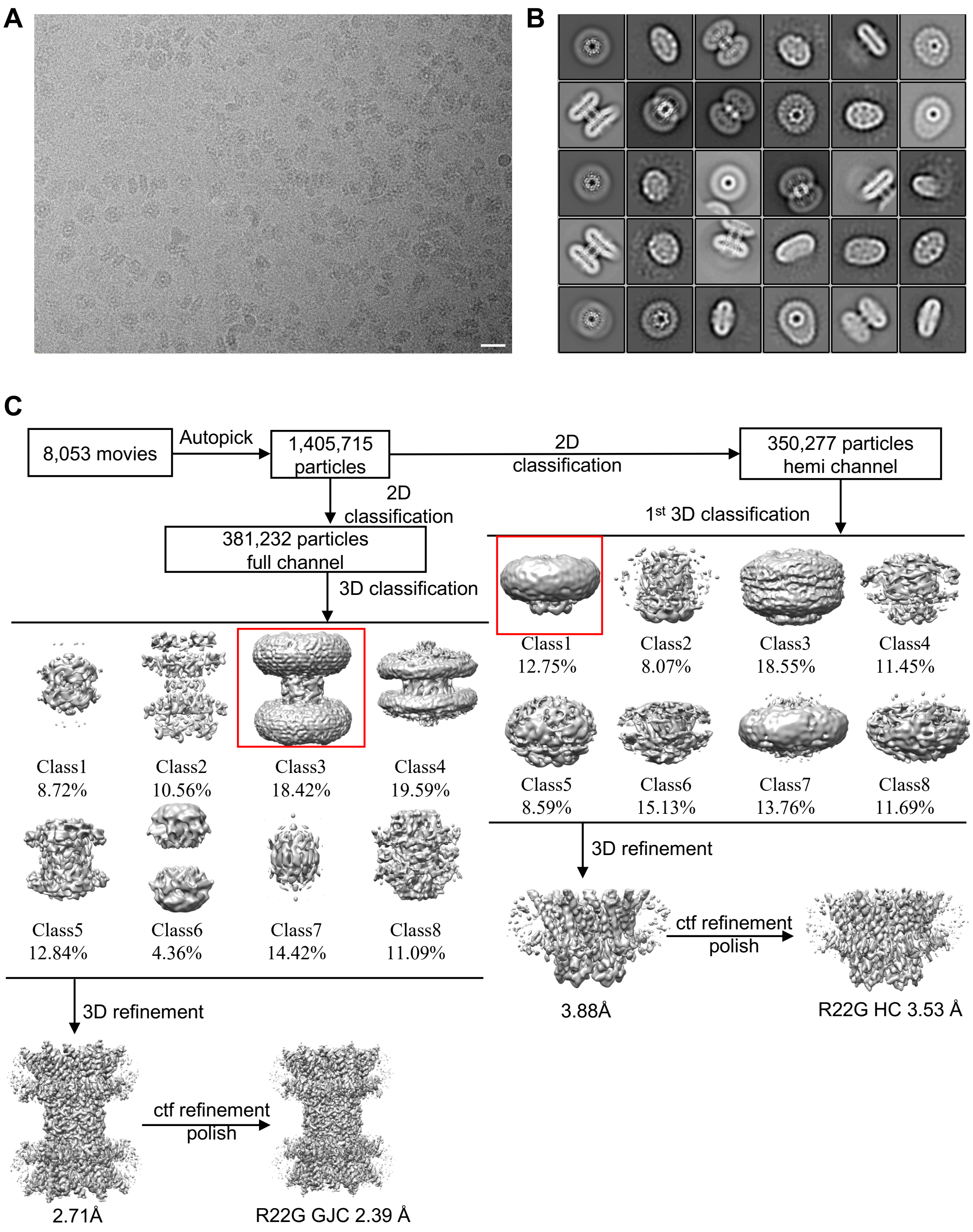


**Figure S5. Cryo-EM image processing pipeline of R22G dataset.** (**A**) A representative micrograph of R22G cryo-EM sample. Scale bar = 200 Å. (**B**) Images of reference-free 2D classification of R22G. The box size is 251 Å. (**C**) Overview of the R22G GJC and HC data processing pipeline.

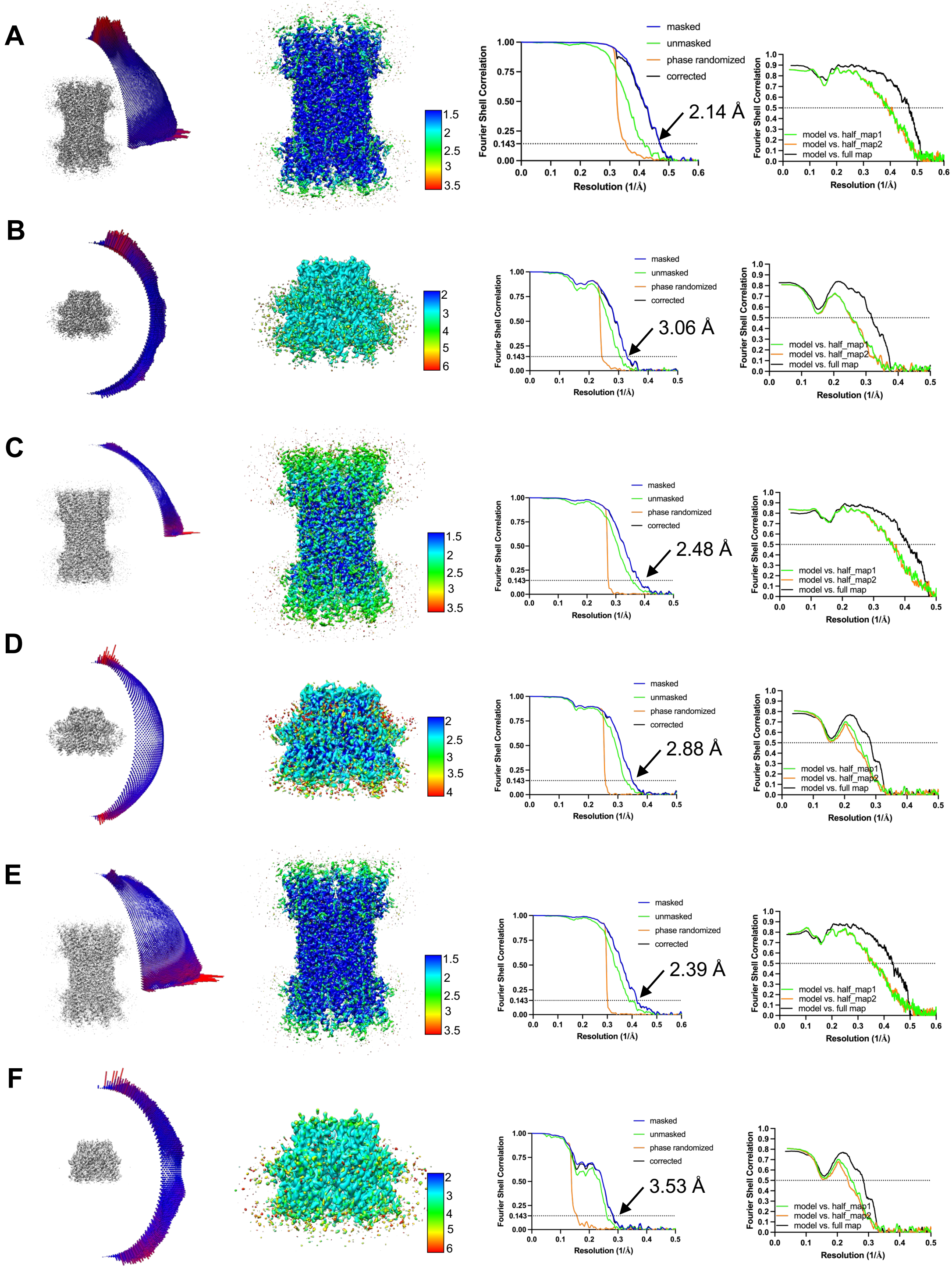


**Figure S6. Angular distribution, local resolution maps and Fourier shell correlation (FSC) plots of wilt-type Cx32 and mutants.** (**A**) Cx32 GJC. (**B**) Cx32 HC. (**C**) W3S GJC. (**D**) W3s HC. (**E**) R22G GJC. (**F**) R22G HC.

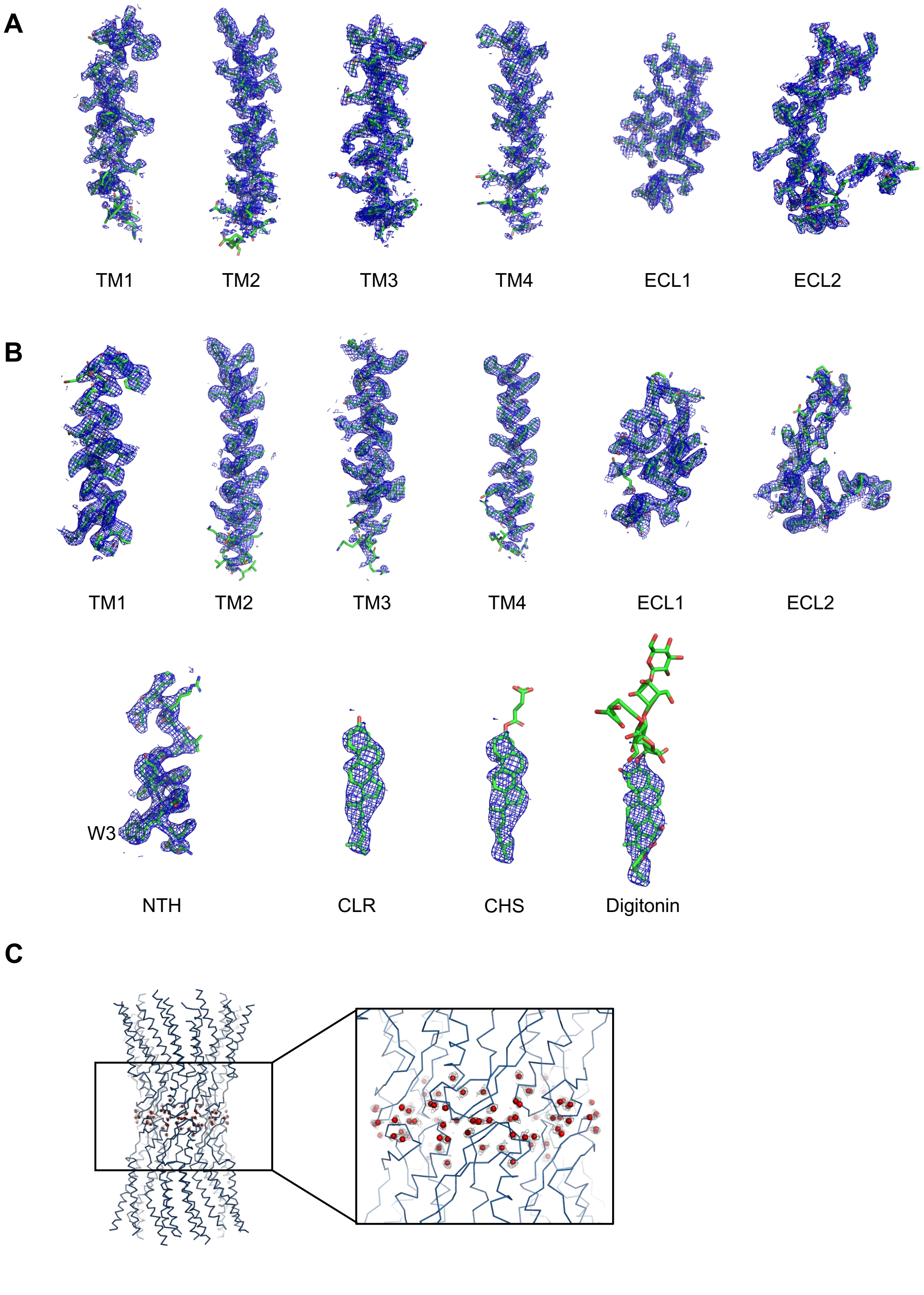


**Figure S7. Representative cryo-EM density map features of Cx32 GJC and HC.** (**A**) Isolated cryo-EM density maps of Cx32 GJC, including the transmembrane (TM) helices and extracellular loop 1 (ECL1) and extracellular loop2 (ECL2)**.** (**B**) Isolated cryo-EM density maps of Cx32 HC, including the TM helices, ECL1, ECL2 and the lipid 2 density under N-terminal helix (NTH) fitting with possible ligands, CLR, CHS and digitonin. (**C**) The observed water molecules in the GJC interface.

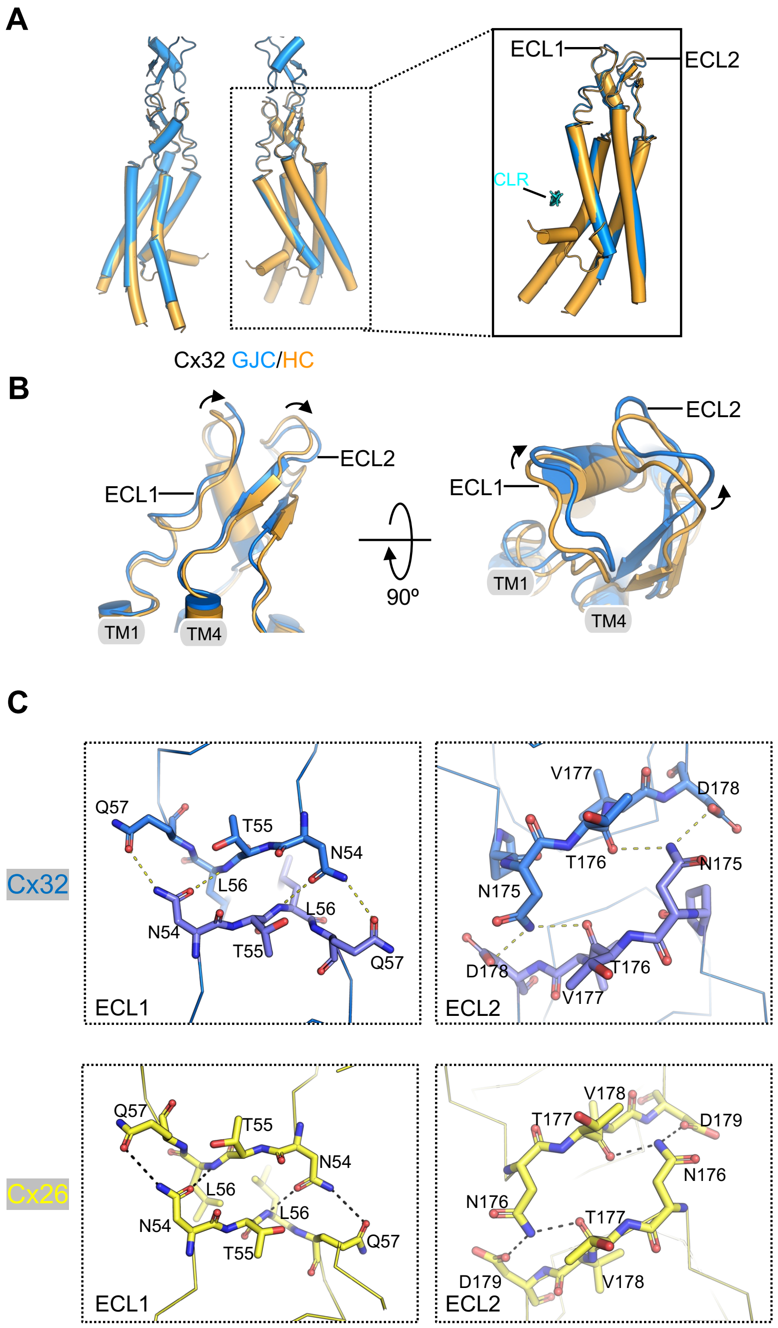


**Figure S8. Structural comparison of Cx32 GJc and HC.** (**A**) Structural alignment of Cx32 GJC and HC. Single subunit alignment of Cx32 GJC and Cx32 HC. The main differences between GJC and HC are the NTH and extracellular loops (ECL1 and ECL2). (**B**) Structural comparison of ECL movement of Cx32 GJC and HC. The arrows indicate the movement of ECL1 and ECL2 of GJC compared to HC. This conformational change was also observed in Cx43 structures. (**C**) Structural comparison of the inter-HC interface. The inter-HC interface of Cx32 (blue) and Cx26 (yellow). The protein sequence of the ECL and structural interaction of Cx32 and Cx26 are similar.

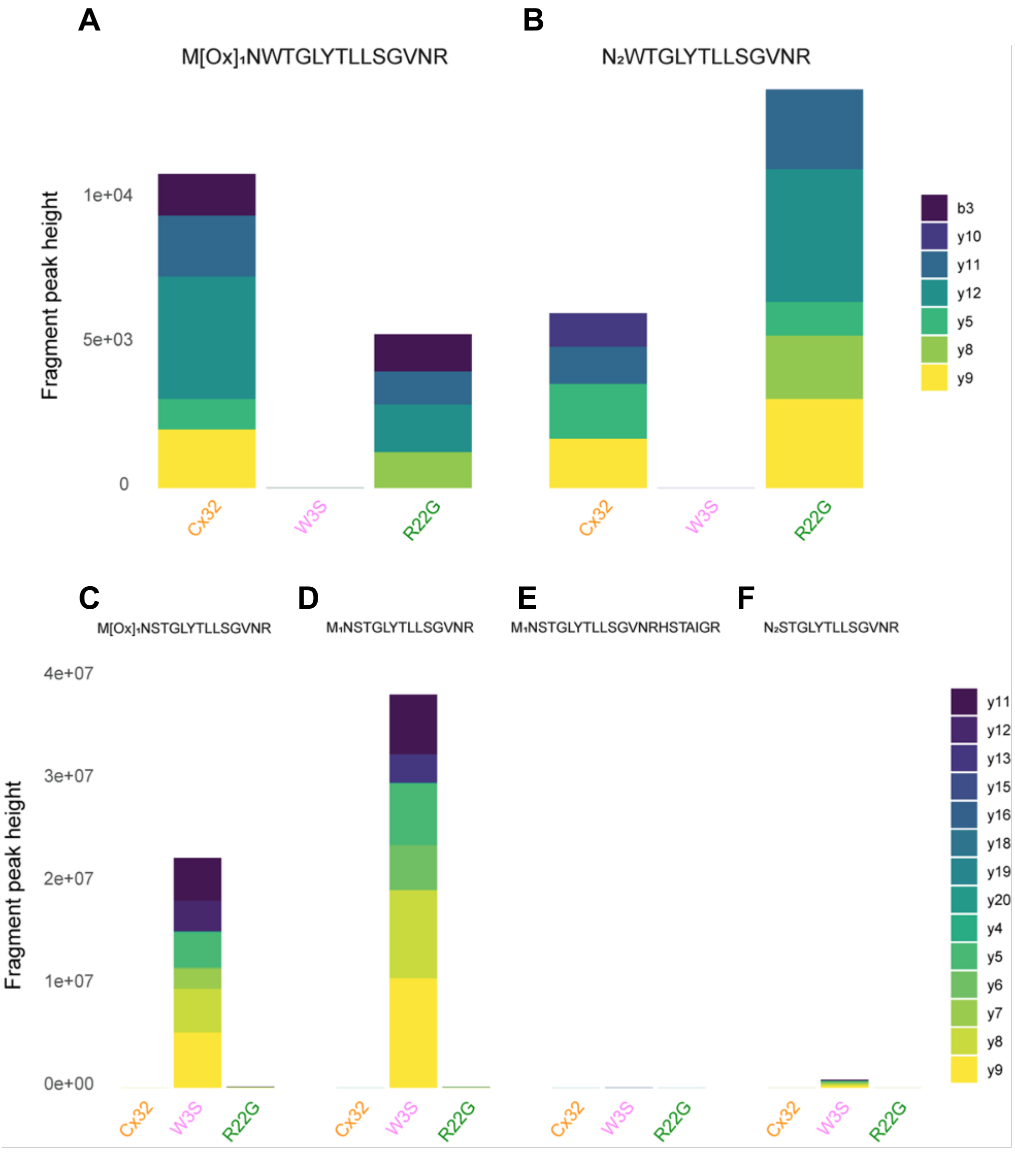


**Figure S9. Mass spectrometric analysis of N-terminal peptides of the purified Cx32.** Analysis of the wild-type Cx32 and the mutants (W3S and R22G) shows that different N-terminal states exist ($M_1$ cleaved and non-cleaved).

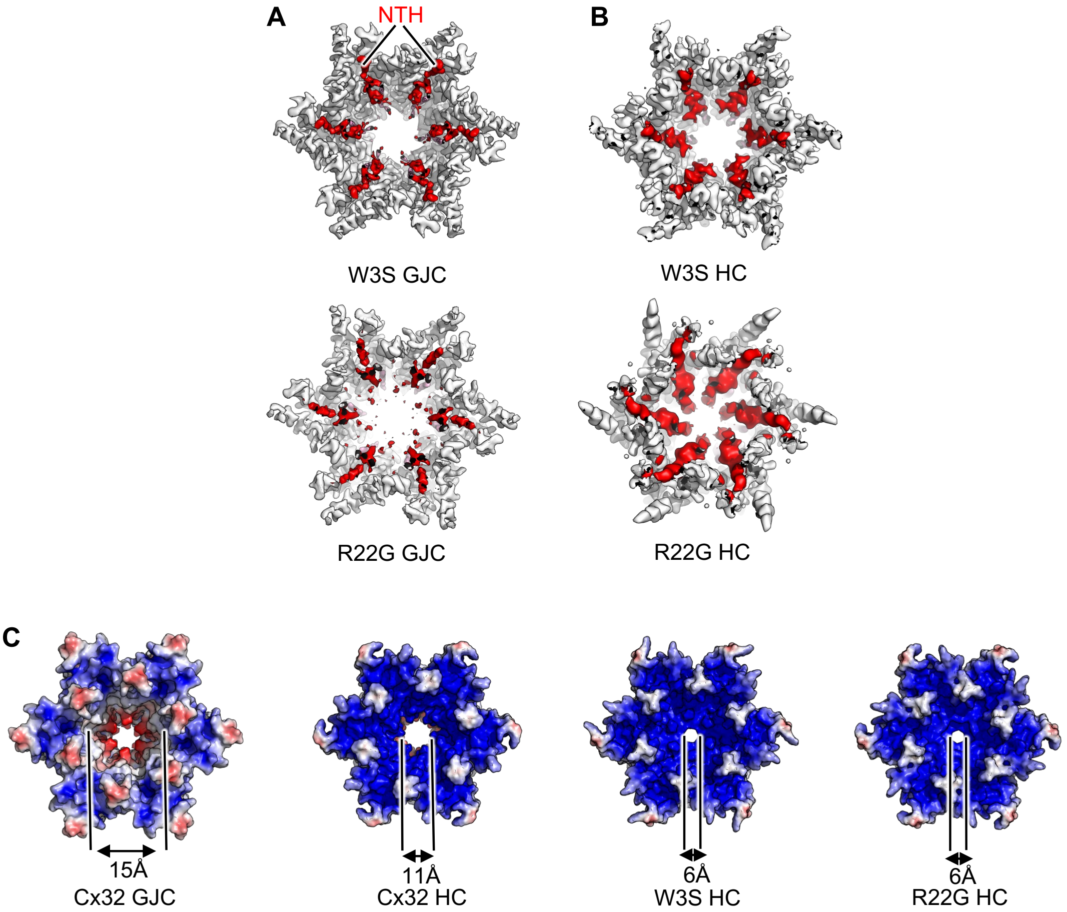


**Extended Figure 10.** (**A**) Cryo-EM maps of W3S and R22G GJCs. The red densities indicate the NTH of W3S and R22G. b, Same as a, for W3S and R22G HCs. (**B**) Surface representation of Cx32, W3S and R22G HCs, colored according to calculated electrostatic potential, show the NTH restricting the connexin HC pore. The diameter of Cx32 GJC is approximately 15 Å, HC is 11 Å, as opposed to 6 Å for W3S and R22G HCs.

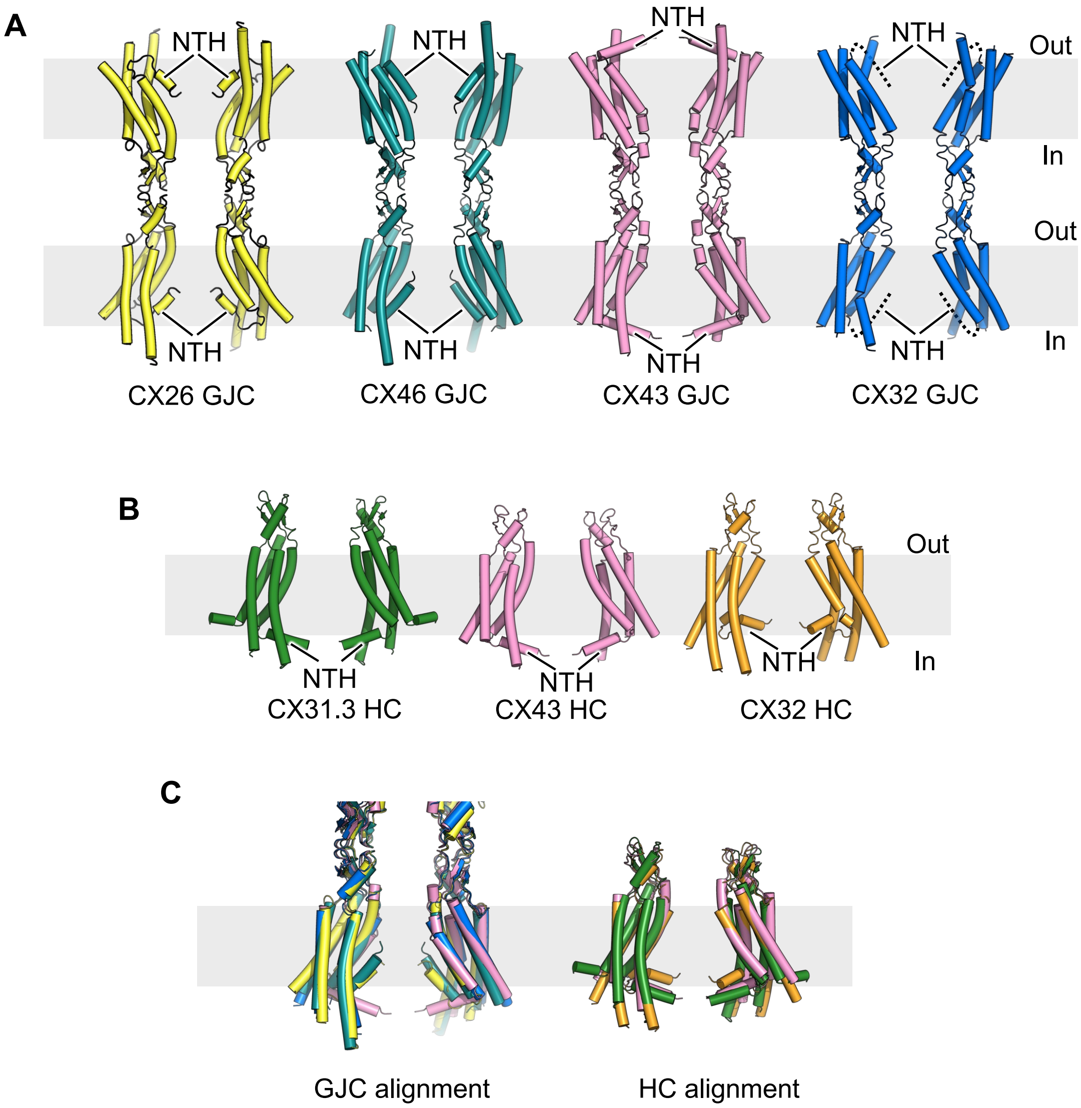


**Figure S11. Structural comparison of the available connexin structures.** (**A**) The structures of Cx26, Cx46, Cx43 and Cx32 GJCs. The NTH of Cx32 is flexible and reminiscent of Cx26 and Cx46. The final model of Cx32 does not include the NTH due to poor cryo-EM density; the dashed line indicates the conformation of the NTH in the Cx32 GJC structure. (**B**) The structures of Cx31.3, Cx43 and Cx32 HCs. The NTH conformation of Cx32 HC is unique, different compared to all other known connexin structures.

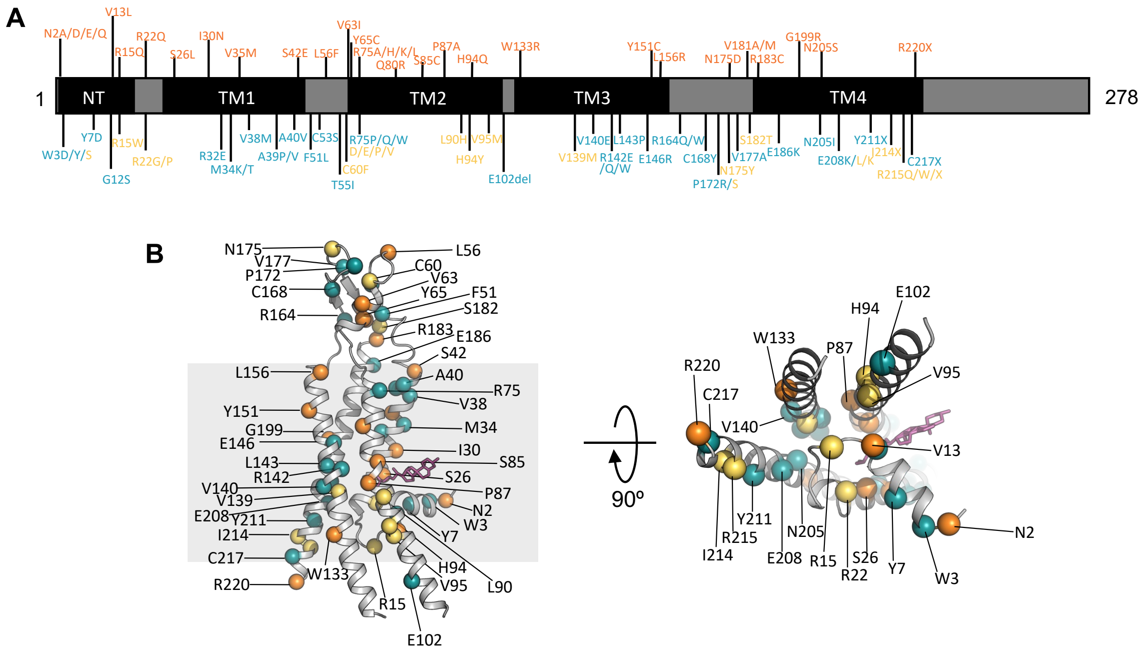


**Extended Figure 12. The Charcot-Marie-Tooth X-linked dominant 1 (CMT1X) disease-linked mutations mapped on the Cx32 sequence and structure.** (**A**) CMT1X mutations are mapped to Cx32 protein sequence. Blue labeled mutations are wrongly assembled and/or not trafficked to plasma membrane; the mutations labeled yellow are able to reach the plasma membrane but do not assemble GJCs; Loss- or gain-of-function mutations are depicted in orange. (**B**) CMT1X disease related mutations are mapped to Cx32 HC subunit structure.

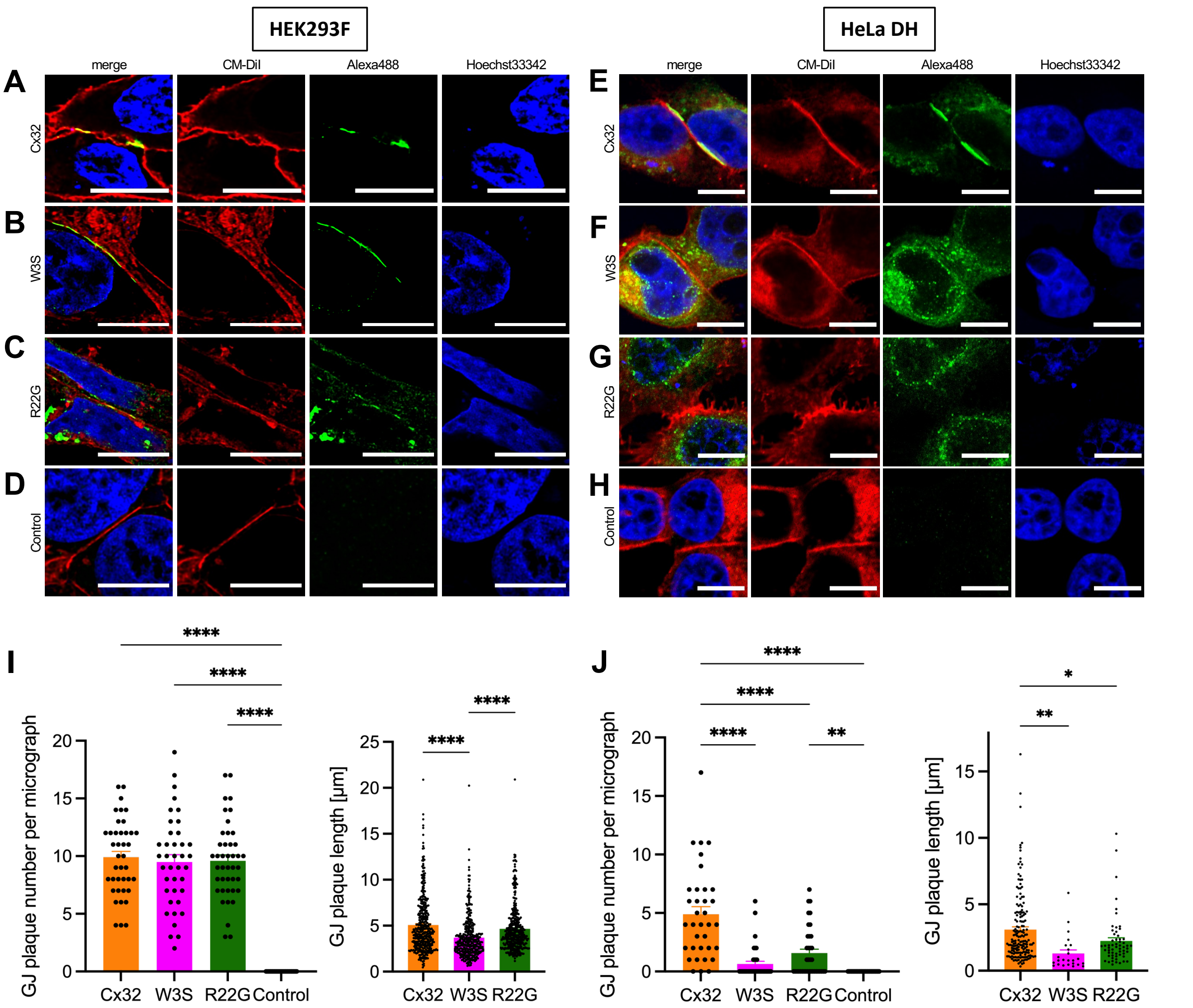


**Figure S13. Immunocytochemistry analysis of Cx32 in HEK293F and HeLa DH cells.** Expression of Cx32, W3S and R22G in HEK293F cells (**A-D**) and in HeLa DH cells (**E-H**) cells assessed by immunocytochemistry (ICC) using rabbit anti-Cx32 primary antibody and goat anti-rabbit Alexa488 antibody (green). (**A-D**) In HEK293F cells, all of the Cx constructs express on the CM-DiI-stained plasma membrane (red) in the form of GJ plaques, visible as lines between two neighbouring cells. (**E-H**) In HeLa DH, Cx32 forms the GJ plaques (e), whereas W3S and R22G mutants show reduced GJ plaque formation (f-h). Nuclei are stained with Hoechst 33342 (blue) to help identify the individual cells. Scale bar = 10 µm. (**I**) In HEK293F cells, the number of GJ plaques per micrograph is not significantly different for wild type Cx32 (n = 40) compared to R22G (n = 42) and W3S (n = 39) mutants. No GJ plaques are present in control sample (n = 49). (**J**) The average length of GJ plaques is not significantly different between wild type Cx32 (n = 396) and R22G (n = 403) mutant, whereas W3S GJ plaques are slightly shorter (n = 370) (****, $P < 0.0001$). (**K**) In HeLa DH, wild type Cx32 (n = 35) forms GJ plaques more frequently than R22G (n = 37) and W3S (n = 37) mutants. There are no GJ plaques in control sample (n = 45). (**L**) Wild type Cx32 (n = 171) GJ plaques are longer than both W3S (n = 24) and R22G (n = 5) plaques (*, $P < 0.05$; **, $P < 0.01$; ****, $P < 0.0001$).

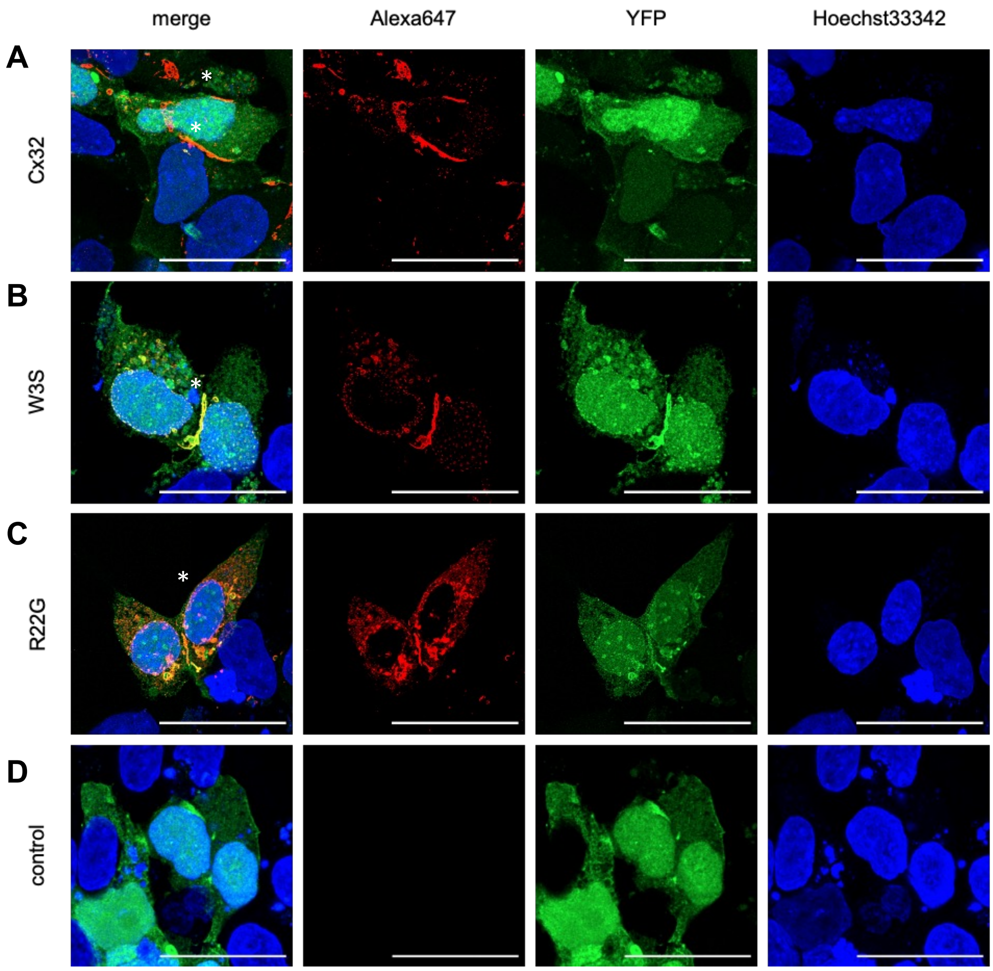


**Figure S14. Transient co-expression of Cx32, W3S and R22G with YFP.** (**A-D**) The images are a maximum-intensity projections of HEK93F cell, co-expressing a test construct with YFP (green): Cx32 (a), W3S (b), R22G (c) or CFP (control, d). The samples were stained using rabbit anti-Cx32 primary and goat anti-rabbit Alexa647 conjugated secondary antibody (red) and Hoechst33342 to stain the nuclei (blue). Asterisks (*) represent the GJC regions, recognized as lines between the cells. Scale bar = 30 µm.

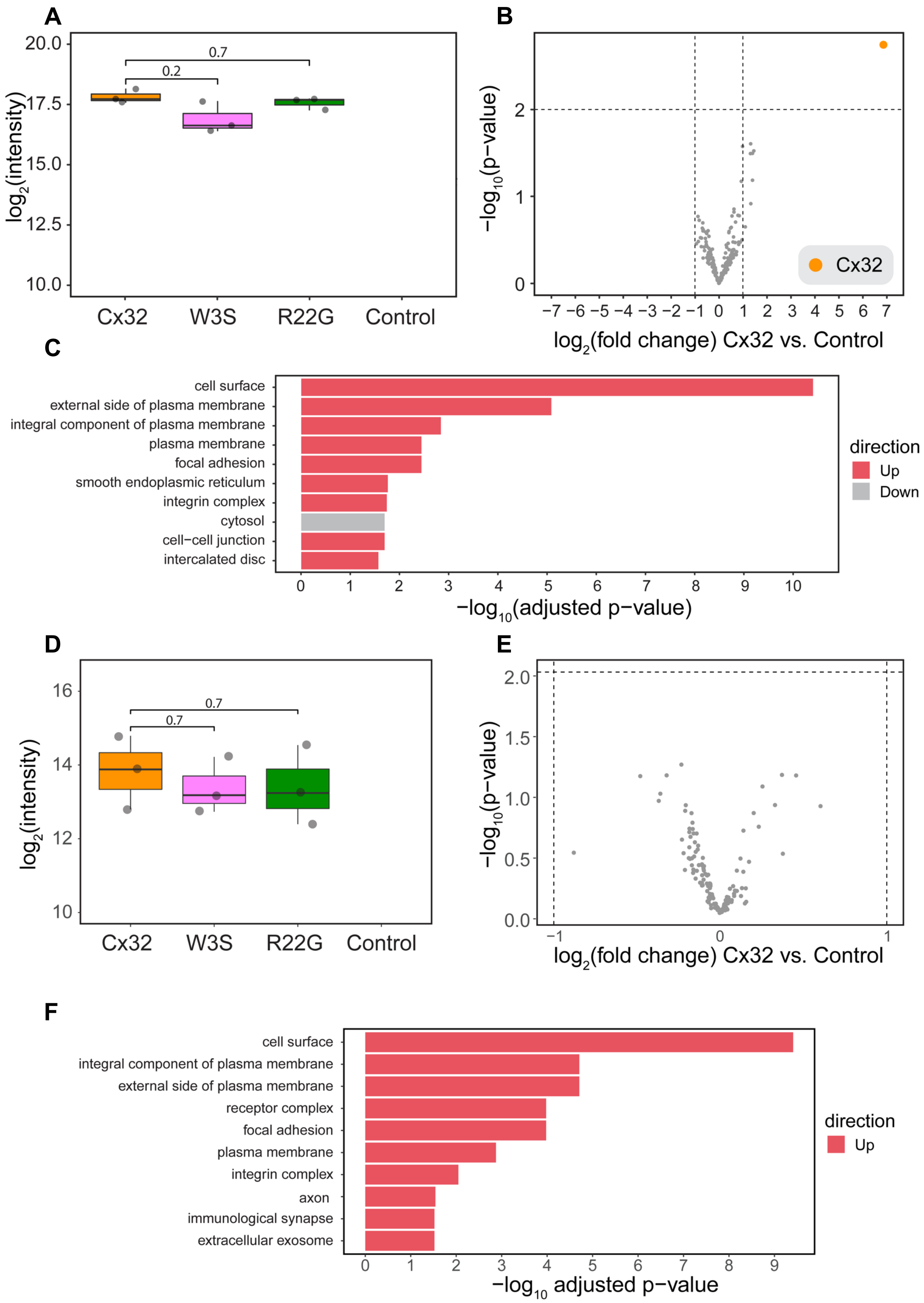


**Figure S15. Analysis of Cx32, W3S and R22G expression by surface protein biotinylation.** (**A**) Mass spectrometric analysis of total Cx32, W3S and R22G expressed in HEK293F reveals that the three proteins express to a similar level in HEK293F cells and that Cx32 is not present in mock-transfected control condition. (**B**) Volcano plot comparing protein abundance of biotinylated proteins identified from the Cx32 WT condition and control. The only significantly changing protein (abs($log_2$(fold change)) > 1 and p-value < 0.01) Cx32 is highlighted in orange. (**C**) GO enrichment analysis of biotinylated proteins shows that they are mainly located on the cell surface. (**D**) Mass spectrometric analysis of total Cx32, W3S and R22G expressed in HeLa cells reveals that the three proteins express to a similar level in HeLa cells and that Cx32 is not present in mock-transfected control condition. (**E**) Volcano plot comparing protein abundance of biotinylated proteins identified from the Cx32 WT condition and control. Cx32 does not show up in this analysis, as CAMthiopropanoylation was not detected on the protein. (**F**) GO enrichment analysis of biotinylated proteins shows that they are mainly located on the cell surface.

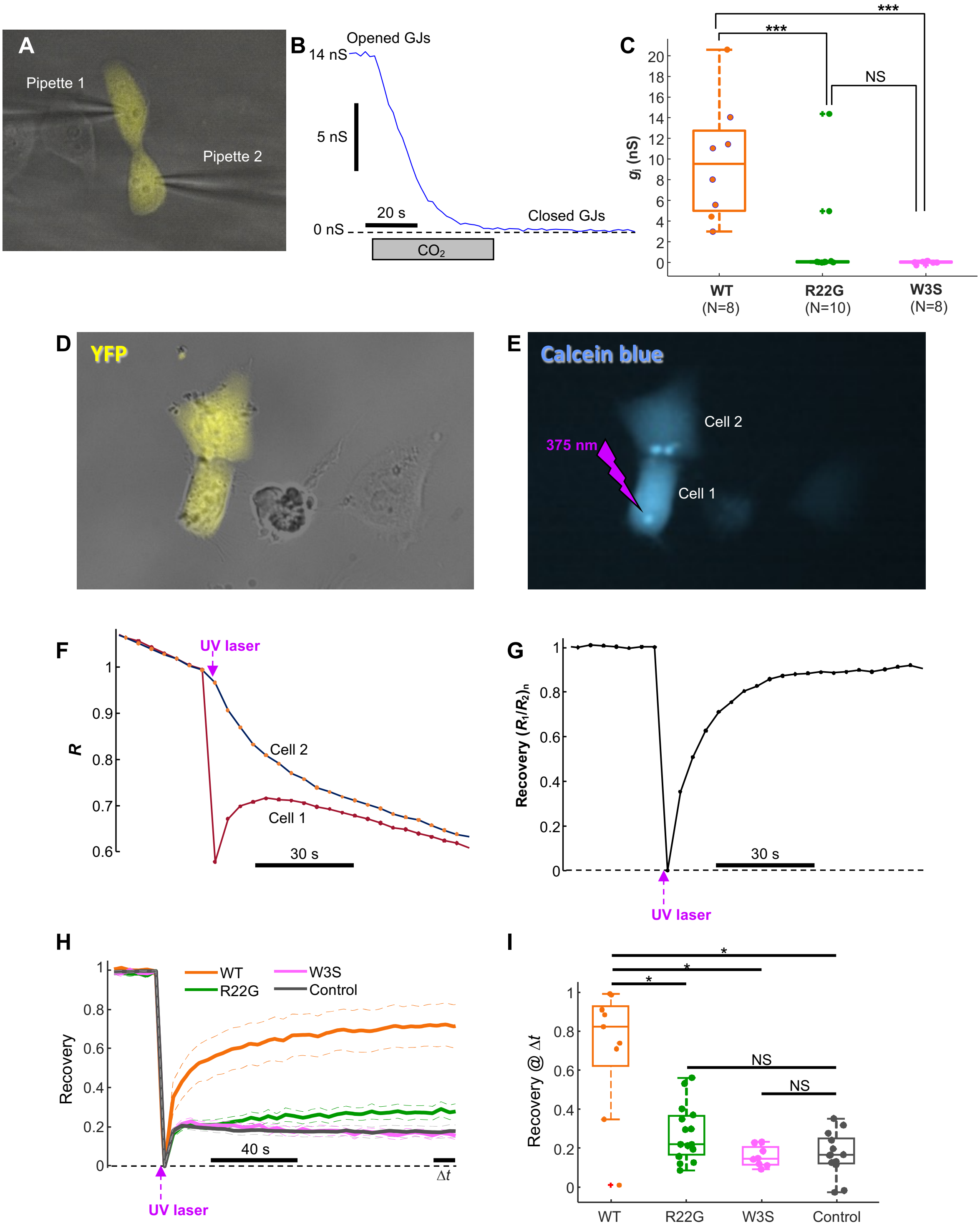


**Figure S16. Permeability properties of WT and mutant R22G and W3S GJCs expressed in HeLa cells.** (**A**) HeLa cell pairs transfected with WT or mutant Cx32 were selected based on their expression of cytosolic YFP and dual patch-clamped to measure the junctional conductance $g_j$. (**B**) $g_j$ was measured in the dual whole-cell configuration while perfusing cells with extracellular solution alone or saturated with $CO_2$ to verify that the junctional current $ij$ was mediated by GJ Channels. (**C**) Box plot of $g_j$ measured in WT (n = 8), R22G (n = 10), and W3S (n = 8) following the same procedure as in panels a-b. In WT experiments, only cell pairs that showed complete uncoupling by the $CO_2$ were retained for the analysis. In the case of the

R22G mutant, two cell pairs have displayed a non-zero junctional conductance, but it was not responsive to the $CO_2$ test, thus we cannot claim that the measured current was mediated by GJs. Statistical analysis was performed using the Kruskal-Wallis test yielding P = 0.001. (**D**) Isolated pairs of Cx32 transfected HeLa cells pre-loaded with Calcein Blue AM and selected for the experiment based on their cytosolic YFP expression. (**E**) A 375 nm UV laser beam is focalized for 300 ms on cell 1 to bleach the Calcein Blue content and observe the following recovery mediated by passive diffusion of Calcein Blue through Cx32 GJs connecting the cell pair. (**F**) Cell 1 and Cell 2 fluorescence traces are normalized as $R = F/F_0$, where $F$ is Calcein Blue fluorescence at time $t$ and $F_0$ is pre-bleaching fluorescence. (**G**) The recovery after photobleaching was quantified as the ratio $(R_1/R_2)_n$ between $R$ in Cell 1 and Cell 2 in a normalized (n) scale between 0 and 1, correspondent to absence of recovery and complete recovery, respectively. This ratio procedure also permitted us to compensate the intrinsic photobleaching of Calcein Blue due to LED excitation (clearly evident in panel f). (**H**) Average recovery after photobleaching for the WT (n = 9), R22G (n = 15) and W3S (n=8) mutations, as well as untransfected (*Mock*) HeLa cells (n=13). Dashed lines are S.E.M. intervals. (**I**) Box plot of individual recovery values computed in the last ten seconds ($\Delta t$) of the experiments averaged in panel h. Statistical analysis was performed by the Kruskal-Wallis test with the post hoc Bonferroni correction, yielding significant (*) differences between the mutants and the WT ($P < 0.05$) and not significant differences with respect to the Mock group.

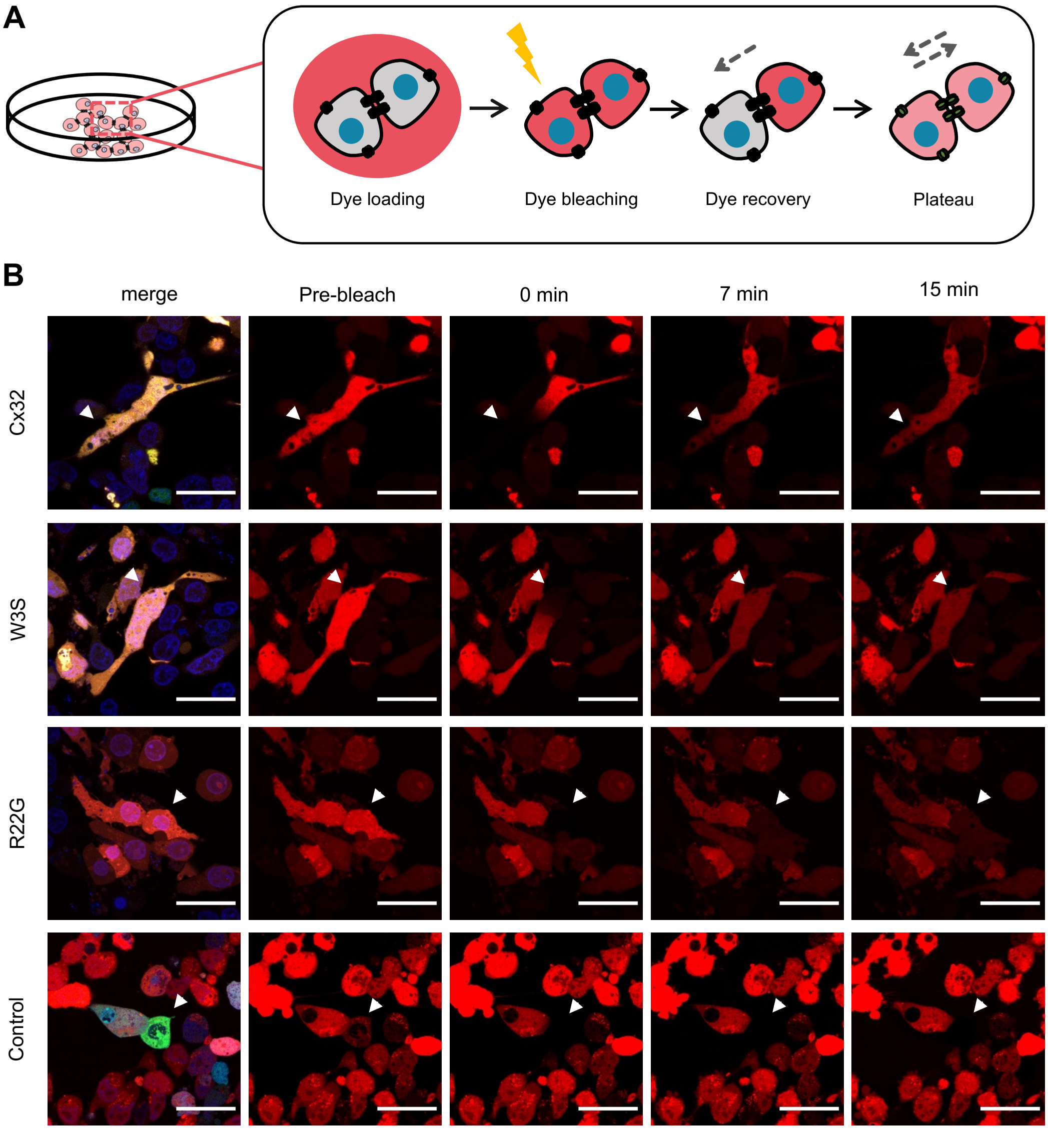


**Figure S17. GJC activity assays in HEK293F cells.** (**A**) Schematic representation of GJC gap-FRAP assay experimental pipeline. The cells are loaded with the dye as in HC dye uptake assays and cells expressing connexin are identified with the microscope. In one cell the dye is bleached and the fluorescence recovery from the neighboring cells, indicative of GJC permeability, observed over time. (**B**) The merged YFP (green), SR101 (red) and Hoechst33342 (blue) channel image was used to select a YFP positive cell, indicative of Cx expression, for bleaching. The cell, which was selected as ROI for bleaching had to be surrounded by neighboring YFP positive cells. The fluorescence recovery was observed until reaching plateau, with representative images shown before bleaching and at 0 min, 7 min and 15 min time points after bleaching. The bleached cell is represented by an arrow. The individual normalized fluorescence recovery curves per sample are shown in graphs next to the respective dye recovery image sequence. Scale bar = 30 µm.

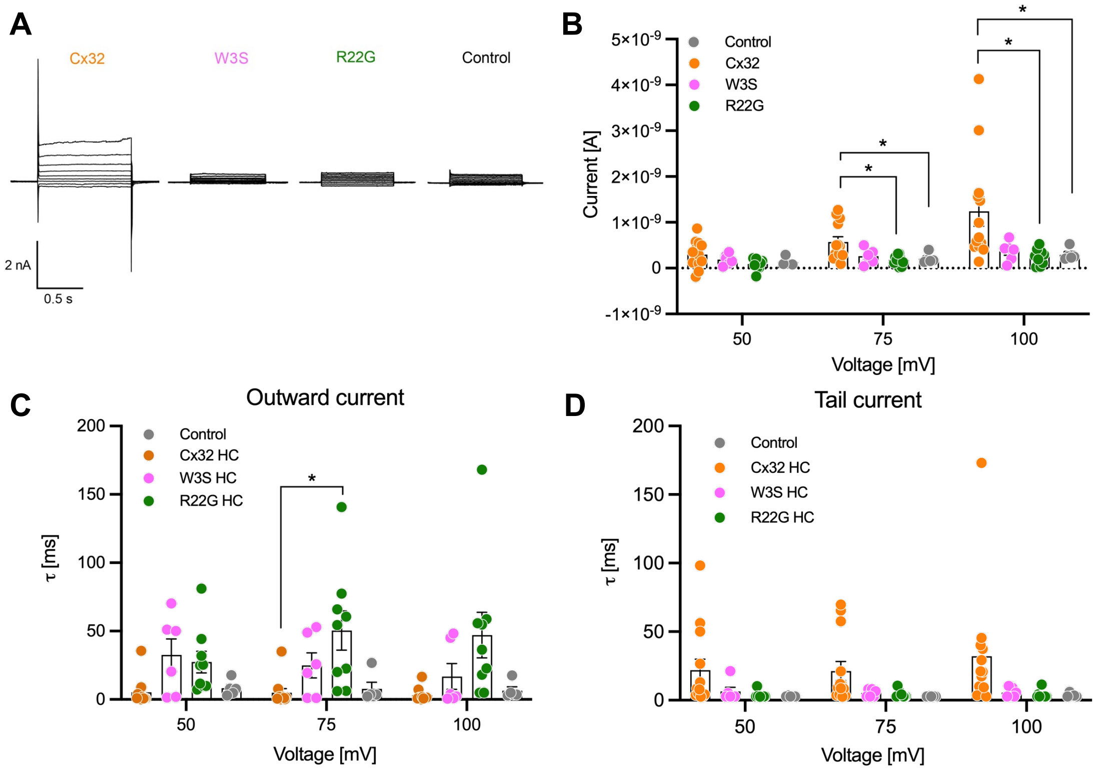


**Figure S18. Biophysical properties of Cx32 hemichannel and mutant currents in HEK293F cells at positive potential of 50 mV, 75 mV and 100 mV.** (**A**) Representative current traces of whole cell patch clamp experiments. (**B**) Peak amplitude of current, Time constant of a single exponential fit for **C,** outward current to steady-state current (n = 5-11) after activation and **D,** tail current (n = 5-13). Data is represented as mean ± SEM and statistical significance was determined by 2-way ANOVA, followed by post *ad-hoc* Dunnett test (*, P < 0.05).

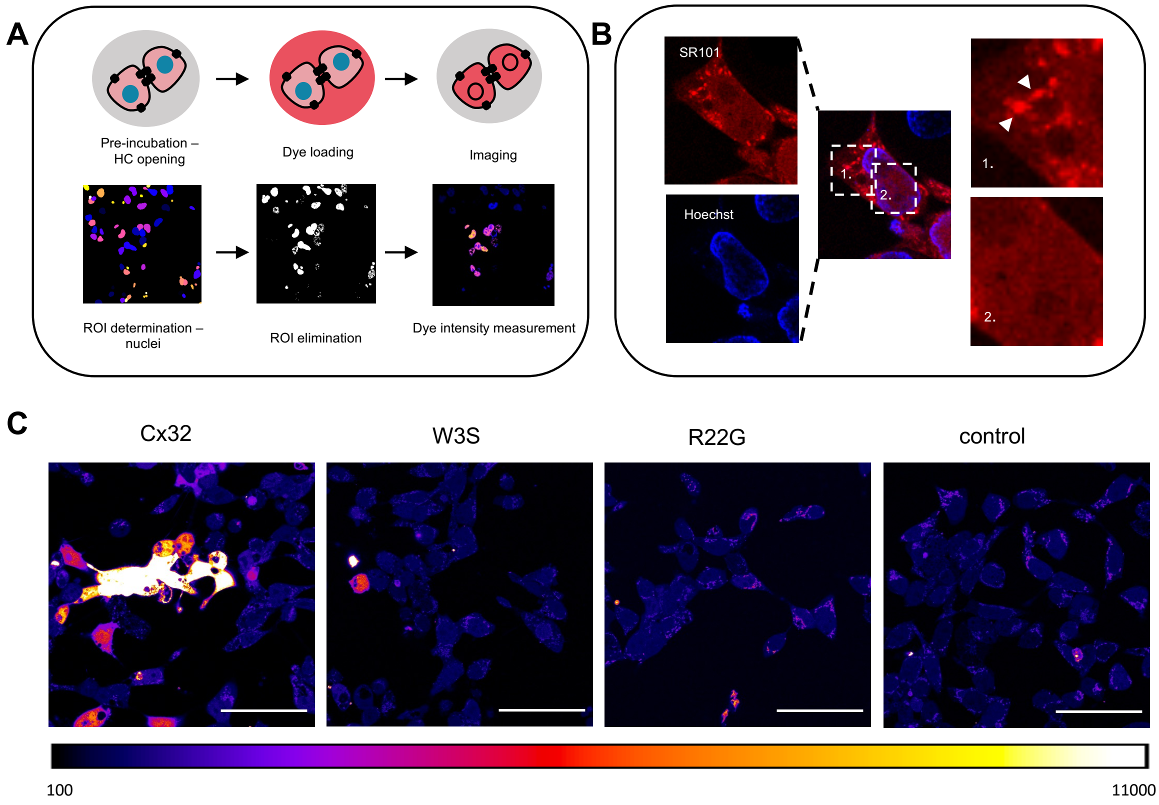


**Figure S19. HC activity assays in HEK293F cells.** (**A**) Schematic representation of HC dye uptake assay experimental pipeline. The cells were pre-incubated in EGTA-containing solution for removal of divalent cations and opening of HCs. The cells were transferred to SR101 and Hoechst 33342 containing solution, washed and imaged. Hoechst 33342 channel images were used for region-of-interest (ROI) determination, YFP channel images for identification of ROIs in transfected Cx-expressing cells, and SR101 channel for intensity measurement. (**B**) Nuclei were selected as ROIs to exclude the unevenly stained cytosolic regions (1.). Since SR101 permeates via HCs and enters the nuclei, which are uniformly stained (2.), the signal measurement leads to higher accuracy. (**C**) The panel shows dye uptake of HEK293F cells transfected with Cx32, W3S, R22G or control. There is a dye uptake increase in Cx32 compared to W3S, R22G, and mock-transfected control. Scale bar = 60 µm.

**Table S1.** Cryo-EM data collection and processing statistics.

| Data collection | | | | | | |
|---|---|---|---|---|---|---|
| Sample | CX32 | | W3S | | R22G | |
| Instrument | FEI Titan Krios/Gatan K3 Summit/Quantum GIF | | | | | |
| Voltage | 300 | | | | | |
| Electron dose($e^-$/Å) | 50 | | | | | |
| Defocus range (μm) | -1 to -2 | | | | | |
| Pixel size (Å) | 0.654 | | | | | |
| Map resolution (Å) FSC threshold 0.143 | CX32 GJC | CX32-HC | W3S GJC | W3S HC | R22G GJC | R22G HC |
| | 2.14 | 3.06 | 2.48 | 2.88 | 2.39 | 3.53 |
| Number of particles | 135,804 | 97,151 | 46,824 | 192,431 | 70,675 | 45,227 |
| Refinement | | | | | | |
| Model resolution (Å) FSC threshold 0.5 | 2.2 | 3.1 | 2.5 | 2.9 | 2.4 | 3.6 |
| Map sharpening b-factor (Å) | -46.41 | -91.18 | -50.67 | -93.48 | -47.69 | -134.135 |
| Map CC | 0.84 | 0.79 | 0.83 | 0.82 | 0.81 | 0.76 |
| Model composition | | | | | | |
| Protein residues/ligand | 2124 | 1242/6 | 2124 | 1236 | 2292 | 1236 |
| ADP (B factor) | 30.08 | 39.33 | 27.69 | 25.34 | 36.68 | 21.74 |
| Bond length r.m.s.d. (Å) | 0.003 | 0.004 | 0.003 | 0.003 | 0.002 | 0.003 |
| Bond length r.m.s.d. (°) | 0.560 | 0.715 | 0.534 | 0.643 | 0.435 | 0.677 |
| Validation | | | | | | |
| MolProbity score | 1.45 | 1.88 | 1.16 | 1.70 | 1.19 | 1.59 |
| Clash score | 3.5 | 11.43 | 2.71 | 7.91 | 3.46 | 7.72 |
| Rotamer outliers (%) | 2.52 | 0 | 1.36 | 0 | 1.17 | 0 |
| Ramachandran plot | | | | | | |
| Favored (%) | 99.42 | 95.57 | 100 | 96.04 | 98.93 | 97.03 |
| Allowed (%) | 0.58 | 4.43 | 0 | 3.47 | 1.07 | 2.97 |
| Disallowed (%) | 0 | 0 | 0 | 0.5 | 0 | 0.50 |

**Movie S1.** The morph of Cx32 compared to W3S.

**Movie S2.** The morph of Cx32 compared to R22G.
